# Aperiodic EEG activity masks the dynamics of neural oscillations during loss of consciousness from propofol

**DOI:** 10.1101/2021.10.12.464109

**Authors:** Niklas Brake, Flavie Duc, Alexander Rokos, Francis Arseneau, Shiva Shahiri, Anmar Khadra, Gilles Plourde

## Abstract

EEGs are known to provide biomarkers for consciousness. Although EEG correlates of loss of consciousness (LOC) are often ascribed to changes in neural synchrony, mounting evidence suggests that some changes result from asynchronous neural activity. By combining EEG recordings of humans undergoing propofol administration with biophysical modelling, we present here a principled decomposition of EEG changes during LOC into synchronous and asynchronous sources. Our results reveal that IPSP decay rate and mean spike rate shape aperiodic EEG features, and that propofol’s effects on these parameters largely explain the changes in EEG spectra following propofol infusion. We further show that traditional spectral EEG analysis likely conflates these effects with changes in rhythmic activity, thereby masking the true dynamics of neural synchrony. We conclude that the well-documented propofol-induced alpha rhythm in fact appears before LOC, and that the moment of LOC is uniquely correlated with the sudden appearance of a delta rhythm.

## Introduction

General anesthesia, a drug-induced state of unconsciousness, is characterized by changes in the EEG. Slow rhythms (< 4 Hz) are a ubiquitous feature of general anesthesia, being readily induced in humans by propofol, sevoflurane, thiopental, and xenon (Guingo et al., 2001; Murphy et al., 2011; Lewis et al., 2012; Purdon et al., 2013; Akeju et al., 2014; Huupponen et al., 2008; Ma and Leung, 2006; Johnson et al., 2003), potentially indicating a universal signature of unconsciousness bridging both general anesthesia and sleep (Amzica and Steriade, 1998; Steriade, 2000; Le Masson et al., 2002). Nonetheless, each anesthetic produces its own characteristic spectral signatures. Most notably, propofol induces a frontal alpha rhythm (8-15 Hz) distinct from any rhythms observed in sleep (Guingo et al., 2001; Feshchenko et al., 2004; Murphy et al., 2011; Ching et al., 2010; Purdon et al., 2013; Akeju et al., 2014). Although it has been suggested that this rhythm plays an important role in propofol’s function as a general anesthetic (Ching et al., 2010, Purdon et al., 2013; Soplata et al., 2017), intracranial LFP recordings indicate that propofol-induced LOC is only coincident with a sharp increase in low-frequency (<3 Hz) power (Lewis et al., 2012). Thus, whether these alpha rhythms play a causal role in propofol’s effects on consciousness remains unclear.

The existence of neural rhythms in EEG is primarily quantified using spectral analysis techniques, such as the spectrogram (Prerau et al., 2016). Peaks in these spectra thus indicate the presence of a neural rhythm (Steriade, 2001; Buzsáki and Draguhn, 2004). However, EEG spectra display not only peaks, but also an overall trend that decays with frequency – variously referred to as the 1/f background, asynchronous, aperiodic, arhythmic, scale-free, or noise component of the signal (Bédard et al., 2006; Miller et al., 2009; He, 2014; Donoghue et al., 2020). It has been recently suggested that this trend may change independently of neural rhythms and that, consequently, changes in frequency-band power over time may not necessarily reflect differences in neural oscillations (Donoghue et al., 2020). Understanding how and why this trend might change is critical to inferring the existence and dynamics of neural rhythms from EEG data.

Recent computational modelling predicted that the spectral trend between 30-50 Hz depends on the average ratio of excitatory and inhibitory synaptic activity, which could be altered by propofol through its potentiation of GABA receptors (Gao et al., 2017). Corroborating this prediction, several recent studies have reported a steeper spectral slope in this frequency range during propofol-induced anesthesia compared to wakefulness (Gao et al., 2017; Colombo et al., 2019; Lendner et al., 2020). These studies have also reported differences in the spectral slope in the 1-40 Hz range (Colombo et al., 2019; Ledner et al., 2020) which, critically, overlaps with the frequencies of propofol-induced neural oscillation, including alpha and delta rhythms. We hypothesized that changes in asynchronous neural activity may act as a confounding variable in traditional analysis techniques when identifying alpha and delta rhythms during propofol-induced LOC.

The goal of this study was to determine how propofol alters the broadband (0.5-100 Hz) aperiodic properties of the EEG, and investigate what implications this may have for interpreting changes in neural rhythms. To this end, we recorded the EEG of 14 human subjects undergoing a fixed-rate infusion of propofol. We then derived a biophysical model of asynchronous neural activity and its influence on spectral EEG properties. This model allowed us to infer changes in the aperiodic components of our EEG recordings simultaneously with changes in neural oscillations. Overall, this study identifies specific physiological parameters that shape the broadband spectral trend in EEG signals, proposes a new method for detrending EEG power spectra, and provides further insight into the mechanisms of propofol-induced LOC.

## Results

### Fixed rate propofol infusion replicates known EEG markers of LOC

To assess neural changes during the transition between wakefulness and unconsciousness, we recorded the EEG of 14 patients during a fixed-rate infusion of propofol (of 1 mg/kg/min) lasting until LOC (Figure 1A). The moment of LOC was identified by the dropping of a held object, which has been shown to provide an accurate, binary measure for LOC (Guay and Plourde, 2019; Cummings et al., 1984). The median LOC-aligned spectrogram was calculated from the Cz channel, revealing the emergence of power within the delta (0.5-3 Hz), alpha (8-15 Hz), and beta (15-30 Hz) frequency ranges (Figure 1B, C). Power in each frequency band started to increase at the same time, but began to decline at distinct times. Beta power began decreasing prior to LOC, while alpha and delta power peaked at or after the moment of LOC (Figure 1C). Similar results were obtained from other electrode locations (Figure S1). These observations suggest that LOC from propofol is associated with the appearance of alpha and delta rhythms, but not beta rhythms, in agreement with past studies (Guingo et al., 2001; McCarthy et al., 2008; Ching et al., 2010; Murphy et al., 2011; Purdon et al., 2013).

**Figure 1.**
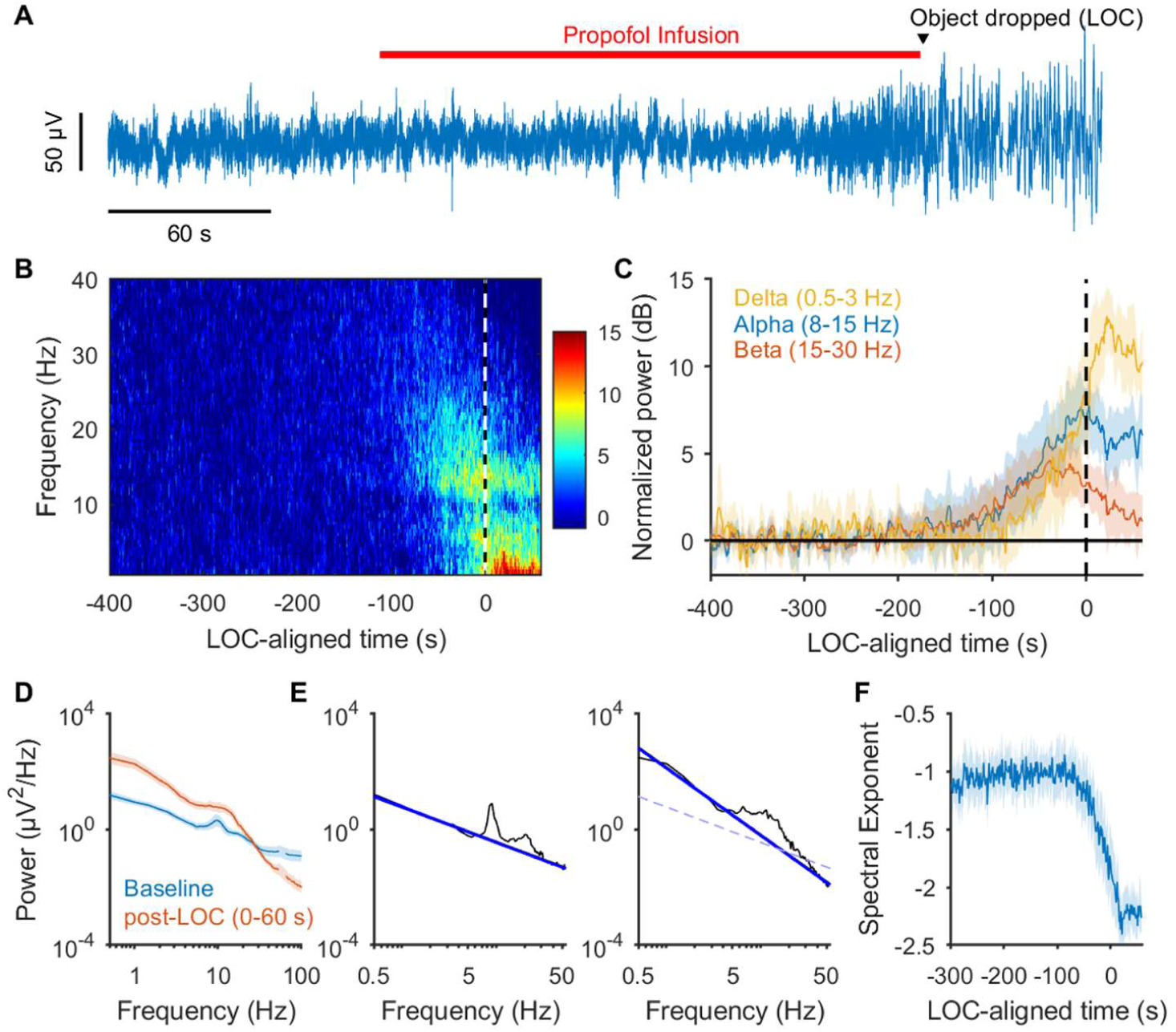
Changes in spectral content caused by propofol infusion. See also figure S1. **(A)**A representative Cz channel recording (Patient 1) shows the appearance of large amplitude, slow oscillations after LOC. The median latency from the onset of propofol infusion to LOC was 135 s (minimum, 95 s; 25th percentile, 109 s; 75th percentile, 177 s; maximum 285 s). **(B)**Average spectrogram from Cz channels of all patients, normalized to baseline power. Color bar ranges from 0-15 dB. **(C)**Time course of delta, alpha, and beta power; shading indicates 95% confidence interval of the mean. **(D)**Average spectra of all patients at baseline (blue) and 0-30 s post-LOC (red). **(E)**Baseline (left) and post-LOC (right) spectrum of Patient 1. Dark blue lines represent fitted 1/f trends between 0.5 and 55 Hz. The light blue dashed line on the right graph reflects the trend at baseline. **(F)**Time course of fitted spectral exponent, i.e. the slope of the spectral trend in log-log space.

To investigate changes in spectral trend, we first compared the power spectra at baseline to that following LOC (0-60 s after the object was dropped). This showed a steepening in the power spectrum around approximately 25 Hz (Figure 1D), consistent with several recent studies (Gao et al., 2017; Colombo et al., 2019; Ledner et al., 2020). To quantify this observation, we estimated the spectral exponent between 0.5 and 55 Hz using the FOOOF algorithm (Donoghue et al. 2020) (Figure 1E). Based on this analysis, we found that that the baseline spectral exponent of -0.93 ± 0.07 (n = 14) decreased to -2.24 ± 0.12 following LOC (p < 10^−6^, paired t-test). To resolve changes in spectral slope over time, we estimated the 1/f trend of the power spectra in 2 second windows; here, we observed a sharp increase in the magnitude of the spectral exponent prior to LOC, followed by a plateau after LOC (Figure 1F). These results corroborate past observations and further demonstrate that changes in the spectral slope are tightly correlated with the moment of LOC.

### 1/f trend does not necessarily reflect asynchronous activity

It has been previously proposed that the 1/f trend present in EEG spectra reflects aperiodic neural activity, while the bumps above this trend reflect power from neural oscillations (Donoghue et al., 2020). Accordingly, fitting a spectral slope may be interpreted as a specific decomposition of the EEG spectrum into periodic and aperiodic power (Donoghue et al., 2020). The forgoing analyses thus provide distinct interpretations of the data; whereas the narrowband analysis indicated a drastic increase in delta rhythms (Figure 1C), the spectral slope analysis classified this power as aperiodic (Figure 1E; Figure 2A-C) and thus did not indicate any significant change in delta rhythms between baseline and post-LOC (Figure 2D). However, this conclusion goes contrary to the inspection of the time series, which clearly showed the appearance of large amplitude, slow rhythmic oscillations following LOC (Figure S2).

**Figure 2.**
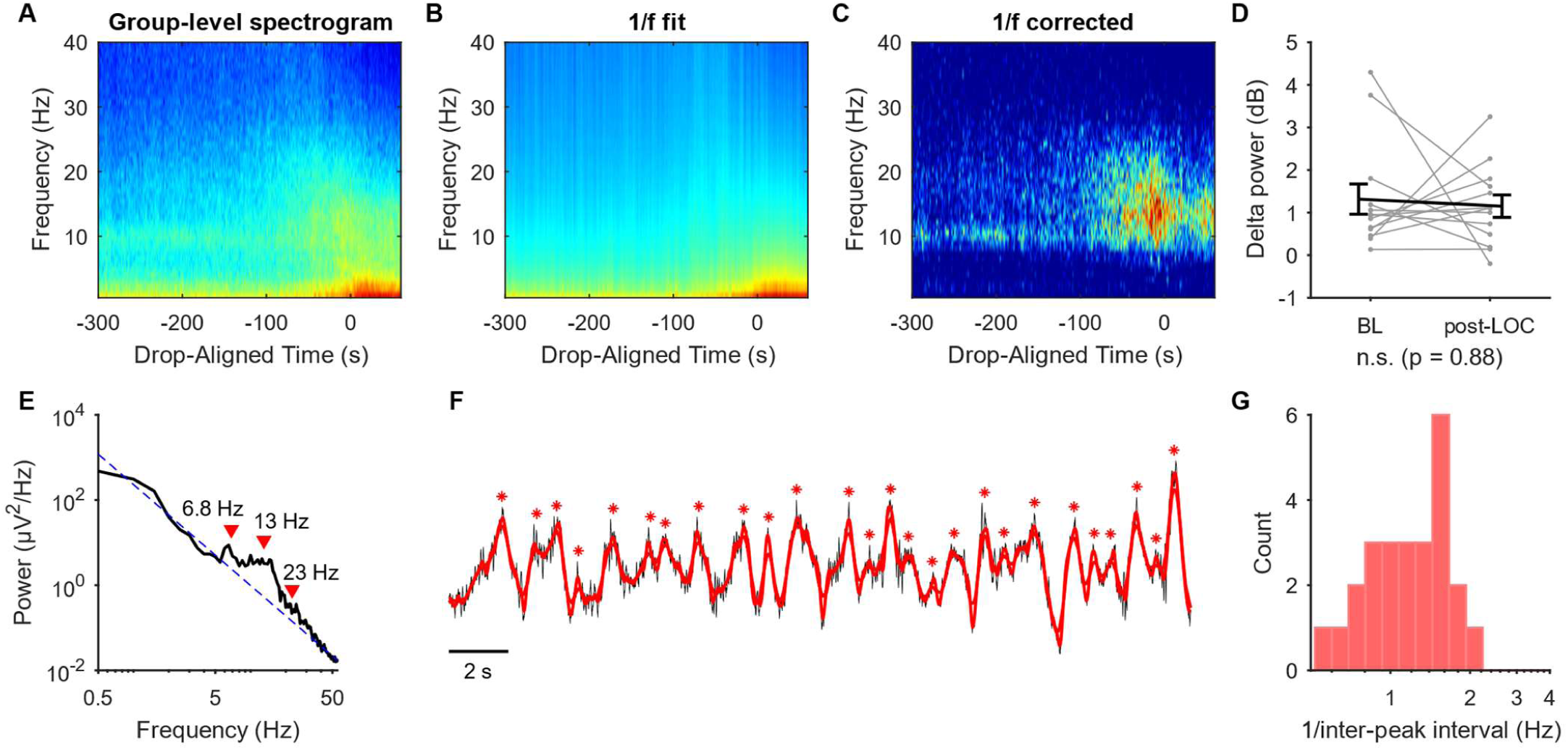
The 1/f trend in EEG spectra conflates rhythmic and aperiodic power. See also figures S2. (**A**) Raw group-level averaged spectrogram. (**B**) Group-averaged 1/f fits. (**C**) Average spectrogram with fitted 1/f background trend removed. (**D**) Delta power, after 1/f trend removed, at baseline (BL) and 0-60 s post-LOC. Grey lines indicate individual subjects, error bars are mean ± SE; a sign test indicates that there is no significant difference change in delta power following LOC (p=0.88). (**E**) An example spectrum post-LOC with a fitted 1/f trend. Peaks at 6.8, Hz, and 23 Hz were identified using the FOOOF package (Donoghue et al., 2020). (**F**) Time series corresponding to the power spectrum shown in panel E (black) as well as the same time series low-pass filtered at 6 Hz (red). Red asterisks indicate peaks in the filtered signal. (**G**) A histogram of the frequency of peaks in panel F, defined as the reciprocal of inter-peak intervals.

Figures 2E-G exemplify the foregoing issue. Figure 2E depicts a power spectrum that was well fit by a 1/f function, indicating no neural oscillations below 6 Hz (Figure 2E). Nevertheless, the corresponding time series displayed slow oscillations (Figure 2F). Algorithmically identifying peaks in the time series after low pass filtering below 6 Hz revealed peaks that occurred with a frequency between 0.5-2 Hz, consistent with the existence of a delta rhythm (Figure 2F,G). It follows that a large fraction of the low frequency power in this spectrum is produced by a delta rhythm, and that the 1/f trend in the spectrum conflates both rhythmic and aperiodic power. Similar analyses of other patients revealed an obvious increase in delta rhythms following LOC in 13 out of 14 patients, while an increase in delta power was only present in 3 out of 14 spectra following 1/f detrending (Figure S2). These results suggested that the empirical 1/f trend is not an appropriate model for aperiodic EEG power, and furthermore questioned whether the aperiodic component does indeed change following propofol infusion, or whether the perceived differences in 1/f trend simply reflects changes in rhythmic oscillations, such as increases in delta rhythms and/or decreases in gamma rhythms (Purdon et al., 2013).

### A model of asynchronous neural activity

To investigate whether propofol changes the aperiodic component of the EEG, we developed a biophysical model for the contributions of asynchronous neural activity to the extracellular electric field. Our goal was to both determine whether it was reasonable to ascribe EEG power to such activity and to derive an analytical expression that could be used to detrend our EEG spectra under specific physiological assumptions.

In Notes S1 and S2, we derived the contributions of asynchronous neural activity to the EEG. Briefly, the model described the expected electric field generated by random, uncorrelated Poissonian synaptic activity, including inhibitory and excitatory post-synaptic potentials (IPSPs and EPSPs) (see Note S1), as well as action potentials (APs) propagating along ascending cortical afferent fibers (Figure 3A) (see Note S2). As a consequence of the linearity of electric fields, we found that under specific assumptions, the EEG signal could be described by the following equation

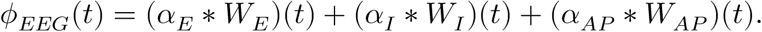

**Figure 3.**
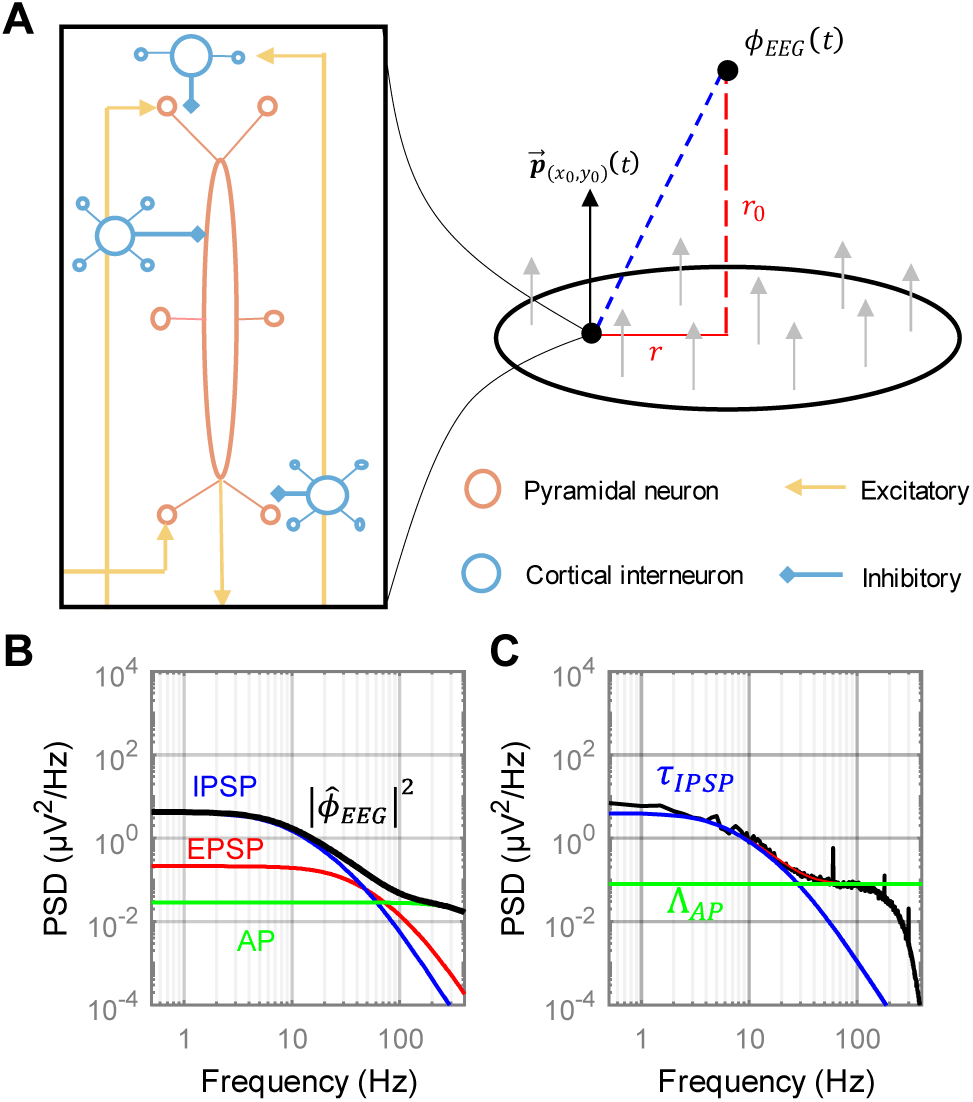
A model of aperiodic EEG power. See also figures S3, S4. (**A**) Left: schematic of local circuitry in neocortex, where each inhibitory (interneuron) and excitatory (pyramidal) cell has been compartmentalized into a single proximal compartment and many terminal compartments. The pyramidal cell compartment is elongated to reflect inclusion of both the soma and apical trunk. Right: the cortex was approximated as a continuum of variously oriented dipole-generating compartments. Modelling synaptic input stochastically allowed the computation of certain properties of the extracellular field generated by asynchronous neural activity. (**B**) The expected power spectral density from aperiodic neural activity determined by the model in Eq. 1 (solid black line), when parameterized using estimates from literature (Table 1). Also plotted are the individual contributions of EPSPs (red), IPSPs (blue) and APs (green). (**C**) The power spectrum of an EEG segment obtained at baseline when no obvious rhythmic activity was observed (black) (see Figure S4). The decay at low frequencies matches the time scale of IPSPs. Although the data is low pass filtered around 300 Hz, a plateau around 100 Hz is visible, as predicted by the model.

In this formalism, *α*_*x*_ represent the responses of the EEG to a single electrical event of each type (*x*= *E, I*, and *AP*), while *W*_*x*_ are stationary Gaussian processes with mean, *μ*_*x*_, and variance, 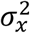. Due to the asynchronicity of synaptic activity, the mean value for *W*_*E*_ and *W*_*I*_ were determined to be 0 (see Note S1). Critically, however, the variances of these Gaussian processes were not 0, and indeed could be calculated as follows

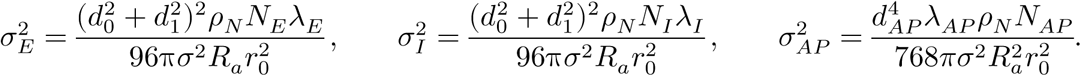

Here, *d*_O_ and *d*_1_ are the average diameters of distal and proximal dendrites, respectively, in the cortex, *ρ*_*N*_ is the density of cells in the cortex, *N*_E_ and *N*_1_ are the total numbers of excitatory and inhibitory synapses per cell in terminal dendrite segments, *λ*_*E*_ and *λ*_1_ are the rates of spontaneous activation for single excitatory and inhibitory synapses, *λ*_*AP*_ is the rate of spontaneous action potential firing along afferent fibers, *N*_*AP*_ are the total number these fibers for each cortical cell, *R*_*a*_ is the axial resistance of cytoplasm, *σ* is the conductivity of the extracellular space, and *r*_O_ is the minimum distance between the electrode and the cortex (see also Table 1).

**Table 1.** Parameters values for model.

| Parameter (units) | Value | Description | Reference |
| --- | --- | --- | --- |
| $r_0$ (mm) | 20 | Scalp-cortex distance | Stokes et al. 2005 |
| $\sigma$ (S/mm) | $5 \times 10^{-6}$ | Volume conductivity | Cosandier-Rimele et al. 2007 |
| $R_a$ ($\Omega$ mm) | 2000 | Axial resistivity | Eyal et al. 2016 |
| $\rho_N$ (cells/mm<sup>2</sup>) | $0.2 \times 10^6$ | Cortical cell density | Markham et al. 2015 |
| $N_E$ (synapses/cell) | 7,500 | No. terminal excitatory synapses | See methods. |
| $\lambda_E$ (Hz) | 0.15 | Spontaneous EPSP rate / synapse | Manns et al. 2004 |
| $\gamma_E$ (μV) | 1000 | EPSP amplitude | Williams & Stuart 2002 |
| $\tau_E^{rise}$ (s) | $1 \times 10^{-3}$ | EPSP rise time | Williams & Stuart 2002 |
| $\tau_E^{decay}$ (s) | $5 \times 10^{-3}$ | EPSP decay time | Williams & Stuart 2002 |
| $d_0$ (mm) | $1 \times 10^{-3}$ | Terminal dendrite diameter | Benavides-Piccione et al. 2020 |
| $d_1$ (mm) | $3.3 \times 10^{-3}$ | Proximal dendrite diameter | Benavides-Piccione et al. 2020 |
| $N_I$ (synapses/cell) | 2,000 | No. terminal inhibitory synapses | See methods. |
| $\lambda_I$ (Hz) | 1.8 | Spontaneous IPSP rate / synapse | Gentet et al. 2012 |
| $\gamma_I$ (μV) | 750 | IPSP amplitude | Larkum et al., 1999 |
| $\tau_I^{rise}$ (s) | $3 \times 10^{-3}$ | IPSP rise time | Thomson et al., 1996 |
| $\tau_I^{decay}$ (s) | $20 \times 10^{-3}$ | IPSP decay time | Thomson et al., 1996 |
| $N_{AP}$ (afferents/cells) | 8,500 | No. (afferent) axons/cell | See methods. |
| $C_m$ (F/mm<sup>2</sup>) | $0.9 \times 10^{-8}$ | Membrane capacitance | Gentet et al., 2000 |
| $R_m$ ($\Omega$ mm<sup>2</sup>) | $0.2 \times 10^6$ | Axonal membrane resistivity | Arancibia-Cárcamo et al. 2017 |
| $\lambda_{AP}$ (Hz) | 0.5 | Spontaneous AP rate | Manns et al. 2004 |
| $d_{AP}$ (mm) | $0.7 \times 10^{-3}$ | Axonal diameter | Arancibia-Cárcamo et al. 2017 |
| $\gamma_{AP}$ (μA/mm) | 0.9 | AP current density | See methods. |
| $\tau_{AP}^1$ (s) | $0.6 \times 10^{-3}$ | Current time constant 1 | See methods. |
| $\tau_{AP}^2$ (s) | $0.4 \times 10^{-3}$ | Current time constant 2 | See methods. |

Because activity was assumed to be uncorrelated, the computed expected power spectrum was found to be

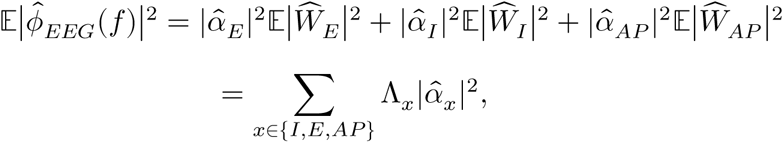

Where 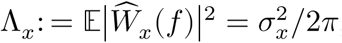, with

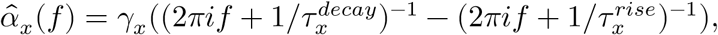

for *x*= *E* and *I*. The equation 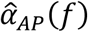in general could not be analytically computed, but the numerically computed power spectrum is shown in Figure S3.

Parameterizing Eq. 1 with values obtained from the literature (Table 1) produced the power spectrum shown in Figure 3B. Interestingly, this parameterization predicted that low frequency power is predominantly produced by inhibitory neurotransmission, while high frequency power is primarily shaped by AP activity (Figure 3B). At lower frequencies, the contributions of IPSPs dominated as a result of their higher frequency of occurrence and slower time scales compared to EPSPs (Table 1); these parameters compensated for the lower density of inhibitory synapses at dendritic tips. On the other end of the spectrum, the fast time scales of APs led to a frequency response largely greater than ∼100 Hz (Figure 3B,Figure S3), and were thus not relevant for understanding aperiodic power in the frequency range of neural oscillations (Figure 3B). Together, these observations suggested that the following reduced model, given by 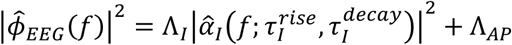, would suffice to capture the aperiodic component of the EEG spectrum. Indeed, this simplified equation succeeded in capturing the spectrum obtained from one of the patients who did not display obvious rhythmic activity on their EEG prior to propofol infusion (Figure 3C; Figure S4).

To understand how aperiodic power interacts with that from synchronous neural oscillations, we returned to our model derivation and, instead of considering purely asynchronous activity, electrical events were made to follow cyclic Poisson processes (see Note S3). Doing so led to an expression identical to Eq. 1, except for the replacement of Λ_*x*_(*x*= *I, E*, and *AP*) by Λ_*x*_(1 + *b g*(*f*)), where *g*(*f*) is a probability density function describing the frequency of fluctuations in firing rate (e.g. Figure 2G) and *b*is a scaling factor related to both the fraction of neurons participating in the rhythm as well as the degree to which the rhythm modulates their firing frequency. This equation extends linearly to Λ_*x*_(1 + ∑_*i*_*b*_*i*_⋅ *g*_*i*_(*f*)) in the presence of multiple modulatory rhythms. Based on this, we can arrive at the following, simplified model of an EEG spectrum:

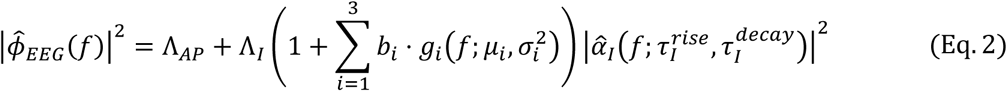

### Model predicts mechanism for spectral steepening following propofol infusion

If the proposed model given by Eq. (2) is valid, we should be able to determine which parameters are responsible for the changes in the EEG power spectra during LOC. To accomplish this, we first fitted the model to the averaged spectra at baseline and at 0-60 s post-LOC. The model fitted the data well (e.g. Figure 4A), and produced estimates for IPSP time constants at baseline consistent with expectations 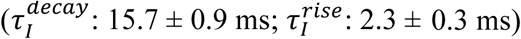. Comparing parameter estimates between baseline and post-LOC revealed significant differences in the parameters 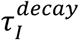, Λ_*I*_, and Λ_*AP*_ (Figure 4B-D). Specifically, 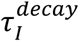increased to 40.0 ± 2.5 ms (n = 13, p < 10^−6^; paired two-tailed t-test), indicating a slowing of IPSP decay of about 150%. Interestingly, this estimate is consistent with experimental observations of propofol’s effect on inhibitory synaptic currents (Orser et al. 1994; Whittington et al., 1996; Kitamura et al., 2013). The scaling factor Λ_*I*_ increased by about 40% (median fold change: 1.37; p = 0.034, sign test), which might suggest an increase in IPSP amplitude; however since both frequency and amplitude affect this scaling factor, it was not possible to discern how much each component was contributing to this change. Finally, Λ_*AP*_ decreased by about 70% (median fold change: 0.29, p < 10^−3^, sign test). This decrease reflects a drop in background cortical input and agrees well with the 50-90% decrease in mean firing rate reported *in vivo* across cortex and thalamus (Redinbaugh et al., 2020; Bastos et al., 2021). Overall, these results validated our predictions and established the model as a plausible framework to characterizing aperiodic EEG power.

**Figure 4.**
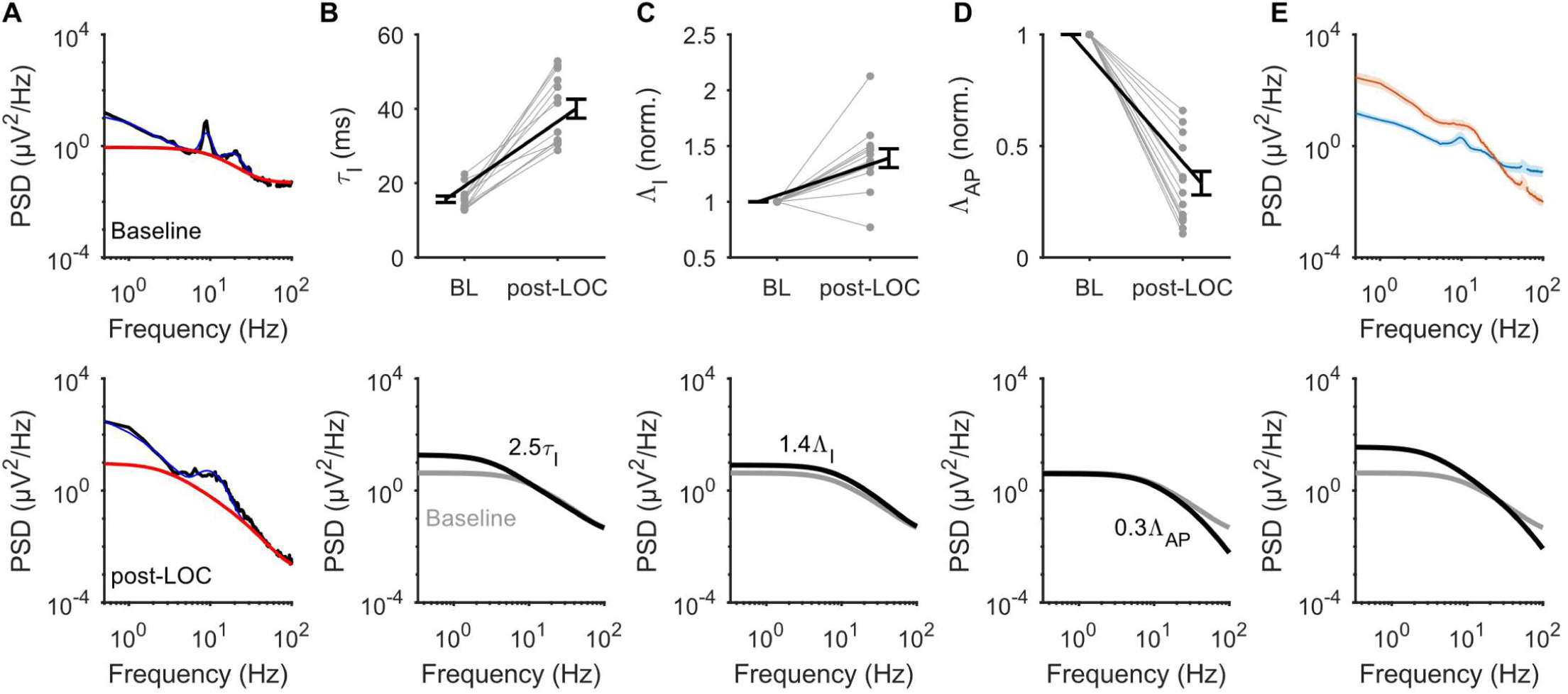
Estimated neural parameters before and after LOC from propofol. See also figure S5 (**A**) Example of model fits to average baseline and post-LOC spectrum of Patient 1. Blue lines reflect the full model fit, whereas the red lines indicate the aperiodic component, i.e. Eq. 2 with *b*_*i*_= 0 for all *i*. (**B**) Top: Estimated decay time of IPSPs 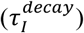at baseline (BL) and post-LOC. Bottom: Grey is the model outcome depicted in Figure 3B. The black line represents the model outcome after increasing the decay time of IPSPs 2.5-fold. (**C**) Same as panel B, but for the scaling factor of IPSPs (Λ_*I*_). (**D**) Same as panels B and C, but for the scaling factor of APs (Λ_*AP*_). (**E**) Top: Average power at baseline (blue) and post-LOC (red). Shading indicates 95% confidence interval of the mean. Bottom: The black line depicts change in modelled aperiodic power after increasing 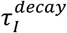2.5-fold and decreasing Λ_*AP*_by 70%.

We next asked what impact the changes in 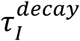, Λ_*I*_, and Λ_*AP*_ should have on the EEG power spectrum. To answer this question, we considered the original model, given by Eq. (1), parameterized with values from the literature (Figure 3B). With this model, we found that slowing the decay of IPSPs by 150% and increasing their amplitude by 40% led to an increase in power at lower frequencies (Figure 4B,C). Meanwhile, decreasing the firing rate of APs by 70% led to a decrease in power across all frequencies, with a stronger effect on higher frequencies (Figure 4D). Modifying the IPSP parameters and the AP firing rate simultaneously led to a rotation of the predicted aperiodic power spectrum in a manner almost identical to that observed experimentally (Figure 4E). Altogether, these results indicated that propofol’s known potentiation of IPSPs and suppression of neuronal spiking are theoretically sufficient to cause a rotation of the EEG power spectrum without any changes to neural synchrony.

### Moment of LOC is uniquely correlated with the appearance of delta rhythms

Our results thus far suggest that not all propofol-induced spectral changes should be ascribed to differences in neural synchrony. Consequently, a spectral analysis that is normalized to baseline power may be systematically misattributing certain spectral changes to differences in synchronous rhythms. Our next goal was to estimate changes in neural oscillations while correcting for anticipated changes in asynchronous neural activity. Fitting our model, given by Eq. (2), to 2 s non-overlapping windows of patients’ spectrograms produced good matches with the data (e.g. Figure 5A). The spectrograms of each patient were then split into periodic and aperiodic components power (Figure 5A). The aperiodic power was defined as the output of the fitted model with the amplitude of oscillations set to zero (i.e. *b*_*i*_= 0, for *i*= 1,2,3). By removing the estimated aperiodic power at each time point from the original power spectrum, we were left with the predicted power from neural rhythms, referred to hereafter as the model-corrected spectrogram.

**Figure 5.**
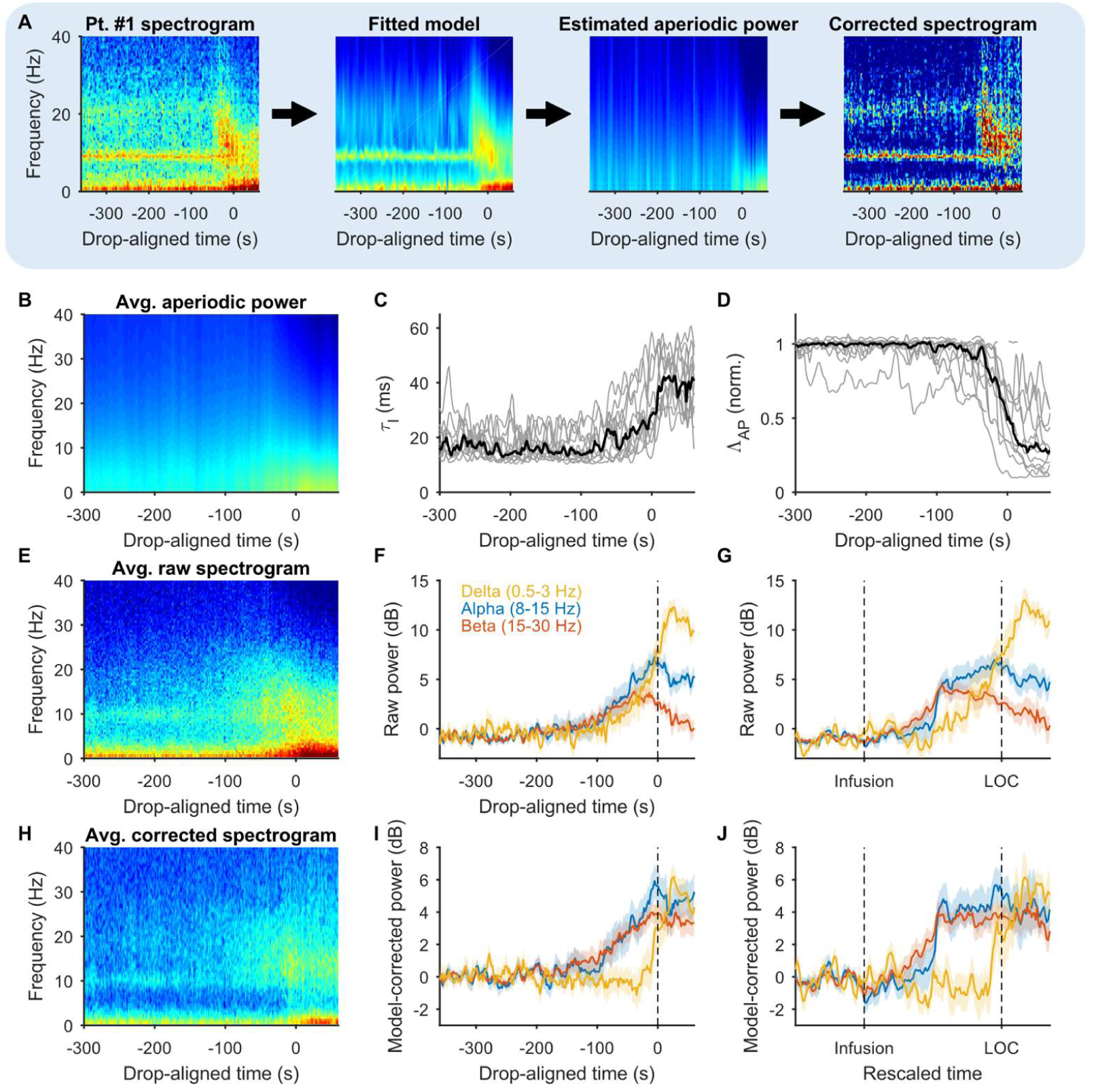
Inferred changes in neural rhythm after aperiodic power correction. (**A**) Example of correction process. From left to right: raw spectrogram from patient 1; output of Eq (2) at each time point after fitting to data; output of Eq (2) with *b*_*i*_= 0 for *i*= 1,2,3, i.e. the aperiodic component of the model; the corrected spectrogram for patient 1, defined as the raw spectrogram divided by the aperiodic power estimated by the model. Note: the colour scale in each individual plot was chosen to maximize contrast, and thus colours are not directly comparable between plots. (**B**) The estimated aperiodic component averaged across all patients. There is a decrease in power above 20 Hz around LOC, indicated by the hue turning darker blue. The lighter hues at low frequencies indicates an increase in low frequency power. Together, this indicates a rotation of the aperiodic component, as seen in Figure 4. (**C**) Time course of the parameter estimates for 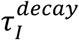 and (**D**) Λ_*AP*_ (**E**) Group-averaged raw power spectrum. (**F**) Time course of power in delta (0.5-3 Hz), alpha (8-15 Hz) and beta (20-30 Hz) bands of the raw spectra; same as Figure 1C. (**G**) Same as in panel F, but plotted against rescaled time so that the time of infusion and LOC are aligned across all patients. (**H**) Group-averaged model-corrected spectrogram. (**I**) Same as in panel F, but for the model-corrected spectra. (**J**) Same as in panel G, but for the model-corrected spectra.

As expected, the group averaged aperiodic power component demonstrated a gradual rotation starting shortly before LOC (Figure 5B). Consistent with our previous results (Figure 4), this temporally-resolved rotation was associated with a gradual increase in 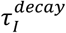and a decrease in Λ_*AP*_ (Figure 5C,D).

We next compared the changes in neural rhythms predicted by the uncorrected, raw spectrogram (Figure 5E-G) and the model-corrected spectrogram (Figure 5H-J). Unlike the raw spectrogram (Figure 5F), the model-corrected spectrogram indicated that alpha, beta, and delta rhythms appeared at distinct times (Figure 5I). Specifically, although both alpha and beta power increased with a similar time course, the increase in delta power was delayed (Figure 5I). Furthermore, whereas the increase in delta power was sudden, the increase in alpha and beta power was gradual as in the uncorrected spectrogram (Figure F, I). We hypothesized that this gradual increase was because the latency from propofol infusion to LOC was variable across patients. To test this, we rescaled time for each patient by dividing LOC-aligned time by the individual latencies to LOC. As expected, following time rescaling, alpha and beta power increased sharply in both the model-corrected and uncorrected spectra (Figure 5G,J). This observation suggests that the appearance of alpha and beta rhythms is more temporally related to the infusion of propofol than the moment of LOC, unlike delta rhythms which were more closely associated with the moment of LOC.

Interestingly, in the raw spectrogram, beta power began decreasing around halfway between propofol infusion and LOC (Figure 5G), eventually producing a beta power not statistically different from baseline 10-30 s post-LOC (p = 0.39, sign test). In contrast, the model-corrected analysis attributes this decrease in the beta band to a broadband decrease in high frequency power caused by a drop in Λ_*AP*_. Consequently, the model indicated that beta rhythms actually remain elevated above baseline following LOC (p < 10^−4^, sign test). Furthermore, in the model-corrected spectra, alpha and beta power appeared linearly correlated throughout the experiment (R^2^= 0.66), which was not apparent in the uncorrected spectra (Figure 5F,G). This correlation suggested that differences in their time courses may be a spurious consequence of aperiodic changes in the EEG.

Finally, as observed with LOC-aligned time, correcting for aperiodic changes had a significant effect on the inferred time course of delta rhythms. In the raw spectra, delta power began increasing gradually around halfway between propofol infusion and LOC (Figure 5G). The model-corrected spectra attributed this slow increase in delta band power to a broadband increase in low frequency power caused by the slowing of IPSP decay. Thus, the model-corrected spectrogram predicts that delta rhythms remain constant until the moment of LOC, at which point they appear suddenly (Figure 5J). In summary, correcting the EEG spectrograms for aperiodic power altered the inferred time courses of neural rhythms, indicating that the alpha/beta rhythm induced by propofol plateaus prior to LOC, while the moment of LOC is uniquely associated with the appearance of delta rhythms.

## Discussion

In the present study, we provided theoretical evidence that propofol leads to aperiodic changes in EEG power spectra at frequencies that overlap with neural oscillations. Our results present a physiological explanation for the broadband EEG changes we observed in humans undergoing LOC from propofol and further demonstrate how these broadband changes alter our understanding of the neural oscillations related to LOC.

### A model for the aperiodic component of the EEG

Generally, identifying oscillations in a noisy signal requires some knowledge of the statistical properties of the noise. The noise in EEG, i.e. the aperiodic component of the signal, has previously been identified with the empirical 1/f trend observed in the power spectrum (Buzsaki and Draguhn, 2004; Demanuele et al., 2007; Donoghue et al., 2020). Although our data displayed an obvious 1/f trend, removing this trend classified all low frequency power as aperiodic, an interpretation of the data which contrasted with the time series behaviour (Figure 2). Indeed, the existence of slow neural oscillations in anaesthetized animals has been widely established, both in EEG and LFP studies, as well as intracellular recordings (Steriade et al., 1993a, b; Steriade, 2001). We concluded that the 1/f trend observed in our power spectra is at least partially supported by high amplitude, low frequency oscillations, and thus conflates power from both periodic and aperiodic neural activity.

Several past studies have postulated physiologically motivated models for the spectral trend that are not purely 1/f (Bédard et al., 2006; Miller et al., 2009; Gao et al., 2017). These studies generally made the same observation that exponential relaxations driven by random spiking display a power spectrum that decays as 1/f above a given frequency, but plateaus at lower frequencies, following a function sometimes called “Lorentzian”, i.e. *P*(*f*) ∝ (1 + *τ f*)^-*β*^. Accordingly, the exponential decays of synaptic currents have been proposed to be a reasonable source of the spectral trend (Bédard et al., 2006; Miller et al., 2009; Gao et al., 2017). The present study concretizes this theory of EEG noise by directly estimating the magnitude of various electrical events to the extracellular field potential. Our results indicated that the overall trend in our data was consistent with these estimates, and furthermore that the effects of propofol observed by others at the molecular and cellular levels are quantitatively compatible with the precise changes we observed in the spectral trend following propofol administration.

Our modelling outcomes contrast with several past studies by attributing low frequency power in the EEG spectrum to inhibitory, rather than excitatory, synaptic transmission (Bédard et al., 2006; Miller et al., 2009). In this same respect, however, our results validated the model of Gao et al. (2017), which assumed a similar baseline ratio of excitation to inhibition to that produced by our model. Nonetheless, our interpretations of the broadband changes caused by propofol differed. While Gao et al. (2017) attributed the steepening of the spectral trend between 30-50 Hz to propofol’s potentiation of IPSC amplitude, this mechanism did not consider the decrease in firing frequency caused by propofol. We found that the combined effect of propofol on frequency and amplitude led only to a net scaling of 140%, which alone was insufficient to cause the increase in low frequency power observed following propofol infusion (Figure 3). Rather, we found that propofol’s effect on IPSC decay contributed the most to altering the power spectrum. Interestingly, because IPSC scaling affected the power spectrum differently than changing the time constant of decay (Figure 4), it is theoretically possible to steepen the spectral trend while maintaining a given excitatory/inhibitory ratio (an example of this is shown in FigureS5). However, we do not contend that this is the case for propofol.

Our model departed from these previous models by including not just synaptic events, but also the explicit contribution of APs. Our results indicated that mean spiking rates were inferable from the EEG signal, as has been previously demonstrated for LFP recordings (Manning et al., 2009; Miller et al., 2009; Buzsaki et al., 2012), although invasive experiments are needed to confirm this. Notably, others have attributed decreases in high frequency power following propofol administration to a decrease in gamma rhythms (Ching et al., 2010; Purdon et al., 2013). The decrease in power we observed was not confined to the gamma range, and in fact extended to the highest frequencies we recorded (Figure 1). We find it is unlikely, therefore, that the parameter Λ_*AP*_ reflects changes in gamma rhythms.

A final theory of EEG noise have posited that the decaying spectral trend arose from low pass filtering effects, such as that from the passive diffusion of synaptic potentials along dendrites (Lindén et al., 2010; Buzsaki et al., 2012). Although we found that the time scales of synaptic activity were sufficient to explain the trend in our data, filtering effects could potentially compound with this trend, leading to steeper spectral slopes than those observed here.

### Neural oscillations related to LOC

Many previous studies of propofol-induced rhythms have not considered changes in aperiodic power when interpreting EEG spectra. Consequently, these studies assumed that all spectral changes reflected differences in neural rhythms. Although correcting for estimated aperiodic changes in the EEG indicated rhythmic power in approximately the same frequency bands as these past studies, it led to important distinctions in the time course of these rhythms.

Increases in delta power have been widely reported during propofol-induced anesthesia (Gugino et al., 2001; Chen et al., 2010; Murphy et al., 2011; Kortelainen et al., 2017). However, the abrupt change observed here has not, to the best of our knowledge, been previously reported in EEG studies, even when the moment of LOC was resolved with high temporal precision (Purdon et al., 2013). We propose that this is because our method of detrending power spectra removes changes in low frequency power attributable to slow changes in IPSP dynamics, which would otherwise be included as oscillatory power. Interestingly, LFP recordings in non-human primates indicated that propofol-induced delta power was tightly correlated with the moment of LOC in the thalamus (Bastos et al., 2021). Thalamic neurons are intrinsically prone to switching from spiking to bursting upon hyperpolarization, a feature that is thought to give rise to the delta rhythm seen in EEG signals (Steriade et al., 1993b; Steriade, 2001). The observed sharp increase in delta rhythms following spectral detrending in this study is consistent with such a binary switching in firing pattern, and suggests that the detrended power spectrum is more consistent with the underlying physiology than the uncorrected spectrogram.

Increases in beta power have been previously observed with low doses of propofol and were associated with the behavioural phenomenon labelled “paradoxical excitability” (Gugino et al., 2001; McCarthy et al., 2008; Ching et al., 2010; Brown et al., 2011). In this study, we also observed an increase in beta power at low concentrations of propofol. Although this increase in beta band power was subsequently followed by a decrease as propofol concentration rose, our analysis suggested that this phenomenon was due to broadband changes in high frequency power caused by decreasing neuronal activity. Whereas past studies have reported a “slowing” of beta rhythms into alpha rhythms (Ching et al., 2010; Purdon et al., 2013), we did not observe this phenomenon after correcting for changes in aperiodic power. Rather, we found that both alpha and beta power appeared at similar times and were tightly correlated throughout LOC. Overall, these results are consistent with the idea that propofol-induced alpha and beta power reflect a single physiological phenomenon (McCarthy et al., 2008; Purdon et al., 2013), but suggest that this phenomenon remains relatively static during LOC.

Frontal alpha rhythms are seemingly unique to anesthetics targeting GABA_A_ receptors (Guingo et al., 2001; Murphy et al., 2011; Purdon et al., 2013; Huupponen et al., 2008; Akeju et al., 2014; Ma and Leung, 2006; Johnson et al., 2003). The emergence of these alpha rhythms has been correlated with the moment of propofol-induced LOC (Ching et al., 2010; Purdon et al., 2013). In contrast, here we observed that the power in this frequency band emerged and plateaued prior to LOC. Notably, we found that this time course was similar in both our model-corrected and baseline-normalized analysis. This inconsistency could be due to the fixed rate of propofol infusion administered here, compared to the target effect-site concentration protocol applied in past studies. Consequently, our analysis of spectral changes pertains to a much shorter time scale of minutes as opposed to hours. Additionally, LOC is marked in this study by the dropping of a held object, unlike past studies that have used auditory stimuli. Our study thus specifically demonstrates that the appearance of alpha rhythms is not an indicator of loss of volitional muscle control. This, however, does not necessarily preclude a role for alpha rhythms in other features of propofol-induced anesthesia.

Overall, our results indicate that there is a clear temporal separation between the appearance of alpha/beta rhythms and delta rhythms during fast induction of LOC with propofol. This observation leads us to conclude that propofol-induced LOC is uniquely associated with the appearance of delta rhythms, consistent with a switching in the thalamo-cortical networks from tonic firing to bursting mode (Steriade, 2001). Our results may indicate that there are sub-anesthetic doses of propofol at which alpha rhythms exists in the absence of delta rhythms. If so, this phenomenon could be used in future studies to isolate and further assess the functional significance of propofol-induced alpha rhythms.

## Methods

### Subjects, EEG recording and anesthetic care

Following MNH Ethics Board approval, we recruited 16 American Society of Anesthesiologists (ASA) class I or II patients (18-65 years old) presenting for lumbar disk surgery as well as 11 control subjects who received no medication. Patients were informed beforehand of the procedures and practiced holding the object. Gold cup electrodes (Fz, Cz, Pz, C3, C4, CP3, CP4, M2 as reference; FC1 as ground; impedance ≤ 5 kOhm) were glued to the scalp to obtain a continuous EEG recording, which was amplified with a 0.1-300 Hz band pass and digitized at 1024 Hz. The standards of care of the Canadian Anesthesiologists’ Society in regard to monitoring, equipment and care provider were rigorously applied. All medications were given intravenously via a catheter placed on the non-dominant arm.

For each patient, we obtained 2 minutes of recording during preoxygenation with eyes closed. The patient was then asked to hold the object (0.5 kg cylinder; 2.5 cm diameter and 15 cm long) in a vertical position with their dominant hand and to keep the eyes closed. The electromyogram (EMG) from the flexor digitorum superficialis and extensor carpi radialis forearm muscle was recorded to monitor the holding ability. Preoxygenation continued for another 2 minutes. Lidocaine 2% (40 mg) was given to attenuate the discomfort caused by the propofol injection. Propofol was given at the rate of 1mg/kg/min and maintained until the cylinder fell from the patient’s hand. Gentle jaw lift was applied if needed to relieve airway obstruction after loss of consciousness. The ability of the patient to respond to loud verbal command was assessed 60 seconds after the fall of the object and the study was then terminated. Two patients were excluded because of failure to comply with the instructions during induction. One patient kept talking, the other kept moving his dominant arm. The final data set was therefore based on 14 patients (10 men; 12 right-handed). Their age and body mass index (BMI) were (mean ± standard deviation) 47.0±11.1 years and 26.5±3.2 kg/m^2^, respectively. They all appeared to be asleep immediately after the fall of the object, with the hand that was holding the object lying immobile. The median duration of the propofol infusion was 135 seconds (range: 95-285), corresponding to a median induction dose of 2.25 mg/kg. Gentle jaw lift was required for 6 patients to relieve upper airway obstruction. Assessment of response to commands 60 seconds after object fall elicited no response or any other form of movement. All 11 controls (6 men) performed the task correctly. Their age and BMI were 22.5±5.2 years and 23.8 ±4.0 kg/m^2^, respectively.

### Data analysis

Spectrograms were computed using the Fast Fourier Transform in 2 second windows with 1.9 s overlap, following the application of a Hamming window. Spectral exponent estimation was done using the Python implementation of the FOOOF package (Donoghue et al. 2020) (FOOOF 1.0.0 executed from Python 3.8.10). All reported values and error bars are given as mean ± S.E. However, shading around lines in plots, e.g. Figure 1D,represent 95% confidence intervals of the mean.

Patient #14 did not display a rotation in their EEG power spectrum following LOC, unlike the other patients. Rather high frequency power increased following LOC. Accordingly, this patient was identified as an outlier by Grubb’s test on the post-LOC Λ_AP_ parameter estimates (fold Λ_AP_ change for patient 14 was 4.02; *c*.*f*. Figure 4D). We thus excluded this patient in the analyses presented in Figures 4 and 5.

### Parameter values

Since excitatory cells are thought to account for 85% of cortical neurons (Markram et a., 2015), we simply used parameters from morphological studies on pyramidal cells to determine the average number of synapses per cortical cell. The number of terminal dendrite segments is equal to 1 more than the number of dendritic branch points, which in L2/L3 neurons has been estimated to be about 50 (Mohan et al., 2015). The average segment length was taken to be about 150 μm (Mohan et al., 2015; Benavides-Piccione et al., 2020; Benavides-Piccione et al., 2021). The values presented in Table 1 for *N*_*E*_and *N*_*I*_ were finally computed by considering a distal excitatory synapse density of 1/μm and distal inhibitory synapse density of 0.2/μm (Karimi et al., 2020).

The rate of spontaneous EPSPs was calculated by multiplying the rate of spontaneous AP firing in excitatory cells (*λ*_*AP*_, Table 1) by a synaptic transmission failure rate of 0.3 (Allen and Stevens, 1994). Similarly, the rate of spontaneous IPSPs was calculated by multiplying the same rate of transmission failure of 0.3 by a spontaneous firing rate for interneurons of 6 Hz (Gentet et al. 2012).

To calculate the number of afferent axons per cells, we used the estimate of 30,000 excitatory synapses per cell (Eyal et al., 2018), and 3.6 synapses/connection (Markram et al., 2015), to arrive at ∼8,500 connections/cell. Finally, the parameters *γ*_*AP*_, 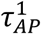, and 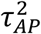were chosen such that simulating an action potential produced a membrane potential waveform with physiologically realistic amplitude and time scale (Figure S3).

### Model fitting

To account for the decay at high frequencies in the data, which likely reflects a combination of high frequency neural noise and amplifier filtering, Eq. (2) was scaled by

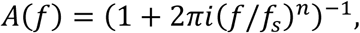

during parameter estimation (i.e. Figure 4 and 5), where *f*_*s*_ and *n* are fitted parameters that did not possess any physiological interpretation. Specifically, the following equation was used to estimate parameters from data

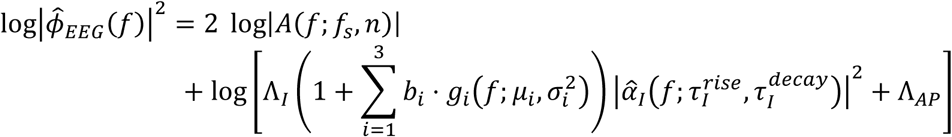

Parameter estimation was performed using a custom written evolutionary-type algorithm with the fitness function

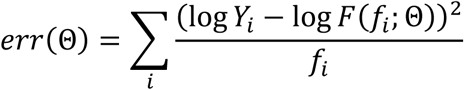

where *Y*_*i*_ is the observed power at frequency *f*_*i*_ (ranging from 0.5 to 250 Hz) and *F*(*f*_*i*_ ; Θ) is the model output for parameter set 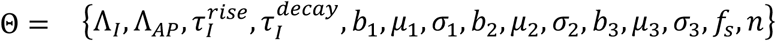. To ensure that the decay at low frequencies was not captured by the function *A*(*f*), the parameter *f*_*s*_ was constrained to be greater than 200 Hz.

The spectrograms were fit in 5 second intervals with 50% overlap, providing parameter estimates for each patient every 2.5 seconds. For the baseline and post-LOC parameters (e.g. Figure 4), the median was taken of these parameters either prior to propofol infusion, or 0-60 seconds following LOC, respectively.

### Code availability

Unless otherwise stated, all data analysis and model implementation was performed in MATLAB 2017b (Mathworks, Natick, MA). All code used for data analysis and figure generation are made available at github.com/niklasbrake/Propofol2021. All data will be made available upon request.

## Authors’ contribution

**Conceptualization:**Niklas Brake.

**Data curation:**Flavie Duc, Alexander Rokos, Francis Arseneau, Shiva Shahiri, Gilles Plourde.

**Formal analysis:**Niklas Brake.

**Funding acquisition:**Anmar Khadra, Gilles Plourde.

**Investigation:**Niklas Brake, Anmar Khadra, Gilles Plourde.

**Methodology:**Niklas Brake.

**Project administration:**Anmar Khadra, Gilles Plourde.

**Resources:**Anmar Khadra, Gilles Plourde.

**Supervision:**Anmar Khadra, Gilles Plourde.

**Validation:**Niklas Brake, Anmar Khadra, Gilles Plourde.

**Visualization:**Niklas Brake.

**Writing – original draft:**Niklas Brake.

**Writing – review & editing:**Niklas Brake, Anmar Khadra, Gilles Plourde.

