## Supplemental Notes for "Aperiodic EEG activity masks the dynamics of neural oscillations during loss of consciousness from propofol"

**Note S1. Contributions of Asynchronous Synaptic Activity to EEG**

To model the EEG signal, we begin by considering the dipoles generated by synaptic activity. Similar to past studies[1,2], we modelled these contributions by considering axial current dipoles in dendrites, given by

$$\boldsymbol{p}_i = I_i L_i \mathbf{e}_i, \qquad (\mathbf{Eq.\,S1}),$$

where $I_i$ is the axial current, $L_i$ is the axial length, and $\mathbf{e}_i$ is the unit vector oriented along the dendrite axis. Previous simulations in pyramidal cortical neurons indicate that synaptic input into the soma and apical trunk produce small dipoles, because these inputs propagate symmetrically, leading to dipoles that largely cancel out[2]. On the other hand, due to the tip effects in more distal compartments, input into these compartments lead to large dipoles[2]. To take this into account, we divide all cortical cells into a single, proximal compartment in addition to smaller distal (terminal) compartments, with the rational that all inputs within the proximal compartment will lead to negligible dipoles. We will assume [**Assumption 1.1**] that these terminal compartments are identical to each other, much smaller than the proximal compartment, and are radially symmetric around the proximal compartment (Figure 2A).

Let us consider such a neuron, with the proximal compartment membrane potential given by $V_0$, and the $N$ terminal compartments given by $V_k,\ k = 1,2,\dots,N$. Because we are considering asynchronous synaptic inputs, almost all voltage changes will be subthreshold. Consequently, we model the neuronal compartments with purely passive membranes, i.e. no voltage-dependent conductances. The membrane voltage in each compartment can then be described by the following system of differential equations[3]

$$C_0 R_0 \frac{dV_0}{dt} = -V_0 + R_0 \sum_{k=1}^{N} \frac{V_k - V_0}{R_{0k}} + R_0 I_0$$

$$C_k R_k \frac{dV_k}{dt} = -V_k + R_k \frac{V_0 - V_k}{R_{0k}} + R_k I_k, \qquad k = 1,2,\dots N.$$

Here, $R_{0k} = R_a L_0 / 2\pi a_0^2 + R_a L_k / 2\pi a_k^2$, where $A = 2\pi a L$ is the surface area of each compartment and $R_a$ is the axial resistance, $C_k = C_m A_k$ and $R_k = R_m / A_k$ are the capacitance and membrane resistance of compartment $k$, respectively. From *Assumption 1.1*, we have that $A_k / A_0 \ll 1$, for all $k \in \{1 \dots N\}$. This assumption effectively decouples the voltage $V_0$ from the terminal compartments. In other words, the voltage fluctuations in the proximal compartment are almost entirely dictated by transmembrane currents according to the equation

$$C_0 dV_0/dt \approx -V_0/R_0 + I_0,$$

whose solution, $V_0(t)$, does not depend on $V_k$. The voltage in the terminal compartments then becomes

$$V_k(t) = \int_0^t \frac{e^{(\tau - t)(1/C_m R_m + 1/C_k R_{0k})}}{R_{k0} C_k} V_0(\tau) d\tau + \int_0^t \frac{e^{(\tau - t)(1/C_k R_k + 1/C_k R_{0k})}}{C_k} I_k(\tau)\, d\tau$$

$$:= V_{0k}^{trans}(t) + V_k^{syn}(t).$$

From Eq. S1 and the fact that dipoles sum linearly, the dipole generated by the entire cell can be describes by

$$\boldsymbol{p}(t) = \sum_{k=1}^{N} \frac{L_0/2 + L_k/2}{R_{0k}} [V_{0k}^{trans}(t) + V_k^{syn}(t) - V_0(t)]\hat{\boldsymbol{e}}_{0k}.$$

It follows from *Assumption 1.1* that $V_{0i}^{trans}(t) = V_{0j}^{trans}(t) = V^{trans}(t)$, $L_i = L_j := L_d$, and $R_{0i} = R_{0j} := R^{trans}$, for all $i, j > 0$, and that $\left(V^{trans}(t) - V_0(t)\right) \sum_{k=1}^{M} \hat{\boldsymbol{e}}_{0k} = 0$ (by radial symmetry). Thus,

$$\boldsymbol{p}(t) = \frac{1}{\widehat{R}_a} \sum_{k=1}^{N} V_k^{syn}(t)\hat{\boldsymbol{e}}_{0k}\,, \qquad (\textbf{Eq. S2}),$$

where $\widehat{R}_a := R^{trans}/(L_0/2 + L_d/2)$. This result is depicted in Figure S3 using a computer simulation of a neuron.

The function $V_k^{syn}(t)$ reflects the voltage changes caused by synaptic inputs, i.e. post-synaptic potentials. In order to simplify *Eq. S2* further, we will assume **[Assumption 1.2]** that $V_k^{syn}(t)$ can be linearly decomposed into the sum of individual post-synaptic potentials. Taking into account both excitatory and inhibitory synapses in a given compartment, *Assumption 1.2* can be formalized as

$$\mathrm{V}_{\mathrm{k}}^{syn}(t) \approx \sum_{j=1}^{N_E'} \sum_{i=1}^{M_E^j} \alpha_E\big(t - E_i^j\big) + \sum_{l=1}^{N_I'} \sum_{m=1}^{M_I^l} \alpha_I(t - I_m^l),$$

where $N_E'$ / $N_I'$ is the number of excitatory / inhibitory synapses in the given compartment, and $\{E_i^j\}_i$ / $\{I_m^l\}_m$ are the times of inputs at excitatory synapse j / inhibitory synapse $l$, which we assume to follow homogenous Poisson point processes with rate $\lambda_E$ / $\lambda_I$. Finally, $\alpha_E$ / $\alpha_I$ are the impulse responses of the membrane potential following excitatory / inhibitory input, both modelled as the difference of exponentials, e.g.

$$\alpha_x(t) = \gamma_x \frac{(\tau_2/\tau_1)^{\tau_2/(\tau_2-\tau_1)}}{1 - \tau_2/\tau_1} \left(e^{-t/\tau_2} - e^{-t/\tau_1}\right)$$

$(x = E, I)$, for $t > 0$, and 0 otherwise. Here, $\tau_1 = \tau_x^{rise}$ and $\tau_2 = \tau_x^{decay}$ determine the characteristic rise and decay times of the EPSP and IPSP, respectively, and $\gamma_x$ determines its magnitude.

Given *Assumptions 1.1* and *1.2*, we can approximate the total dipole moment generated by a single cortical cell by the equation

$$\boldsymbol{p}(t) = \frac{1}{\widehat{R}_a} \left( \sum_{j=1}^{N_E} \sum_{i=1}^{\mathrm{M}_{\mathrm{E}}^j} \alpha_E\big(t - E_i^j\big)\boldsymbol{e}_j + \sum_{l=1}^{N_I} \sum_{k=1}^{M_I^l} \alpha_I(t - I_k^l)\boldsymbol{e}_l \right), \qquad (\textbf{Eq. S3})$$

where $N_E$ and $N_I$ are the total number of excitatory and inhibitory synapses in terminal compartments, and $\boldsymbol{e}_j$ and $\boldsymbol{e}_l$ are the unit vectors oriented from the synapse towards the proximal compartment.

To model the EEG signal, we consider the cortex as a continuous plane and compute the electric field generated at a point $\boldsymbol{x}_0$ (Figure 2A); without loss of generality, we can define $\boldsymbol{x}_0 = (0,0,r_0)$. By modelling all points within an infinite homogenous volume conductor, the electric field may be computed as[4]

$$\phi_{syn}(t) = \frac{1}{4\pi\sigma\widehat{R}_a}\sum_i \frac{\langle \boldsymbol{p_i}(t), \boldsymbol{x}_0 - \boldsymbol{x}_i\rangle}{||\boldsymbol{x}_0 - \boldsymbol{x}_i||^3} \ ,$$

where $i$ sums over all cortical cells, located at positions $\boldsymbol{x}_i$. By accounting for both excitatory and inhibitory cells, the latter equation can be further expanded into

$$\begin{aligned}\phi_{syn}(t) &= \frac{1}{4\pi\sigma\widehat{R}_a}\sum_i \frac{\langle \boldsymbol{p_i}^E(t), \boldsymbol{x}_0 - \boldsymbol{x}_i\rangle}{||\boldsymbol{x}_0 - \boldsymbol{x}_i||^3} + \frac{1}{4\pi\sigma}\sum_i \frac{\langle \boldsymbol{p_i}^I(t), \boldsymbol{x}_0 - \boldsymbol{x}_i\rangle}{||\boldsymbol{x}_0 - \boldsymbol{x}_i||^3}. \\ &\coloneqq \phi_E(t) + \phi_I(t).\end{aligned}$$

Since $\phi_E(t)$ and $\phi_I(t)$ are computed using the same steps, we will only show here the derivation for $\phi_E(t)$.

Let the $i^{th}$ dipole at time $t$ be generated at a distance $r_i$ from the electrode in the cortical plane, with an angle $\phi_i$ relative to the synapse-electrode axis. The equation for $\phi_E(t)$ then becomes

$$\phi_E(t) = \frac{1}{4\pi\sigma\widehat{R}_a}\left(\sum_i \frac{\alpha_E(t - t_i)\sin\varphi_i}{r_0^2 + r_i^2}\right), \qquad (\mathbf{Eq.\,S4}).$$

We model this system using a Poisson point process defined on the space $\{(t_i, r_i, \varphi_i) \in \mathbb{R}^+ \times \mathbb{R}^+ \times [-\pi/2, \pi/2]\}$. By applying change of variables, the density of cortical cells along a circle of radius $r$ is given by $2\pi r\rho_N$, for some area density $\rho_N$. The density of terminal synapses is thus $\rho_N N_E$. Because of our assumption of radial symmetry (*Assumption 1.1*), the density of synapses at a given radius, $r$, and a given angle to the electrode, $\varphi$, becomes $\pi r\rho_N N_E \lambda_E \cos\varphi$. We can thus rewrite Eq. S4 as a sum over discretized space, given by

$$\begin{aligned}\phi_E(t) &= \frac{1}{4\pi\sigma\widehat{R}_a}\sum_{0\le k_1\le N_1}\sum_{0\le k_2\le N_2}\left(\alpha_E * W_{k_1,k_2}\right)(t)\frac{\sin(-\pi/2 + \pi k_2/N_2)}{r_0^2 + (k_1\Delta r)^2} \\ &= \alpha_E(t) * \sum_{0\le k_1\le N_1}\sum_{0\le k_2\le N_2} W_{k_1,k_2}(t)\frac{\sin(-\pi/2 + \pi k_2/N_2)}{4\pi\sigma\widehat{R}_a(r_0^2 + (k_1\Delta r)^2)}\end{aligned}$$

where $W_{k_1,k_2}(t) \sim \text{Poisson}(\pi(k_1\Delta r)\rho_N N_E \lambda_E \cos(-\pi/2 + \pi k_2/N_2)\,\Delta\varphi\Delta r)$.

Let $y_{k_1,k_2} \coloneqq W_{k_1,k_2}(t)\frac{\sin(-\pi/2+\pi k_2/N_2)}{4\pi\sigma\widehat{R}_a(r_0^2+(k_1\Delta r)^2)}$. Since $y_{k_1,k_2}$ are independent under the assumption of asynchronous activity, it follows from the Lyapunov's Central Limit Theorem that

$$\sum_{0\le k_1\le N_1}\sum_{0\le k_2\le N_2} y_{k_1,k_2} \to \mathcal{N}\left(\sum_{0\le k_1\le N_1}\sum_{0\le k_2\le N_2}\mu_{k_1,k_2}\,, \sum_{0\le k_1\le N_1}\sum_{0\le k_2\le N_2}\sigma_{k_1,k_2}^2\right),$$

as $N_1, N_2 \to \infty$. Furthermore, these sums become Riemann integrals as follows

$$\sum_{0\le k_1\le N_1}\sum_{0\le k_2\le N_2}\mu_{k_1,k_2}$$
$$=\sum_{k_2}\sum_{k_1}\frac{\pi(k_1\Delta r)\rho_N\Delta r\lambda_E N_E\sin(-\pi/2+\pi k_2/N_2)\cos(-\pi/2+\pi k_2/N_2)\,\Delta\varphi}{4\pi\sigma(r_0^2+(k_1\Delta r)^2)}$$
$$\to\int_{-\pi/2}^{\pi/2}\int_0^\infty\frac{\pi r\rho_N\lambda_E N_E\cos\varphi\sin\varphi}{(r_0^2+r^2)}drd\varphi=0$$

since $\sin\varphi\cos\varphi$ is an odd function. Similarly,

$$\sum_{0\le k_1\le N_1}\sum_{0\le k_2\le N_2}\sigma^2_{k_1,k_2}\to\int_{-\pi/2}^{\pi/2}\int_0^\infty\frac{\pi r\rho_N\lambda_E N_E\sin^2\varphi\cos\varphi}{16\pi^2\sigma^2\widehat{R}_a^2(r_0^2+r^2)^2}drd\varphi=\frac{\rho_N N_E\lambda_E}{96\pi\sigma^2\widehat{R}_a^2 r_0^2}$$

We thus arrive at the following equation for the EEG signal contributions from EPSPs

$$\phi_E(t)=(\alpha_E\star W_E)(t),$$

where $W_E(t)\sim\mathcal{N}(\mu_E,\sigma_E^2)$, and

$$\mathrm{\mu_E}=0,\qquad\sigma_E^2=\frac{\rho_N N_E\lambda_E}{96\pi\sigma^2\widehat{R}_a^2 r_0^2}.$$

Note that because of the assumed radial symmetry, the expected value is zero as the dipoles will on average cancel out. Critically, however, the variance is not zero, which will play an important role in shaping the aperiodic spectral component of the EEG.

**Note S2. Contributions of Asynchronous Action Potentials (APs) to EEG**

Action potentials (APs) propagating along axons can contribute to the extracellular field potential[5] and are thought to shape the broadband spectral features of LFP recordings[6]. To investigate the impact of APs on the aperiodic component of the EEG, we model here the contributions of asynchronous AP activity to the extracellular field potential.

Let us consider an axon that rises vertically along the apical-ventral axis, before turning parallel to the cortical surface in a manner similar to what have been observed in layer 1 of cortex (Figure 3A). Based on this set up, we can parametrize the axon domain as follows

$$\boldsymbol{g}(l; x_0, y_0, \varphi, L) = \begin{cases} (x_0 + (l-L)\cos\varphi\,, y_0 + (l-L)\sin\varphi\,, L), & l \geq L \\ (x_0, y_0, l), & l \leq L \end{cases}$$

where $(x_0, y_0)$ is the position of the ascending component in the cortical plane, and $L$ is the z coordinate at which the bend occurs. As in the case of synaptic activity, we want to understand the electrical field at a point $\boldsymbol{r} = (0{,}0, r_0)$. Let $I_a(l,t)$ be the axial current at time $t$ and at a position $l$ along the axon. The equation for the electric field is then given by[4]

$$\phi_{AP}(t; x_0, y_0, \varphi, L) = \int_g \frac{\boldsymbol{g}(l) - \boldsymbol{r}}{4\pi\sigma|\boldsymbol{g}(l) - \boldsymbol{r}|^3} \cdot I_a(l,t)\mathrm{d}g(l)$$

$$= \frac{1}{4\pi\sigma}\left[\int_{-\infty}^{L} \frac{I_a(l,t)(r_0 - l)}{(x_0^2 + y_0^2 + (r_0 - l)^2)^{3/2}}\ \mathrm{d}l \right.$$

$$\left. + \int_L^{\infty} \frac{I_a(l,t)(x_0\cos\varphi + y_0\sin\varphi + l - L)}{(x_0^2 + y_0^2 + 2x_0(l-L)\cos\varphi + 2y_0(l-L)\sin\varphi + (l-L)^2 + (r_0 - L)^2)^{3/2}}\ \mathrm{d}l\right].$$

Along an infinite cable, the voltage is given by (Ermentrout and Terman, 2010)

$$V(l,t) = \int_{-\infty}^{\infty} G(l-x,t)V_0(x)dx + \frac{r_M}{\tau}\int_0^t\int_{-\infty}^{\infty} G(l-x, t-s\ )I(x,s)dx\,ds,$$

where $V_0(l)$ is the initial distribution of voltage along the axon, $I(l,t)$ is the transmembrane current in space and time, and

$$G(l,t) = \frac{1}{\sqrt{4\pi\lambda^2 t/\tau}}\exp\left(-\frac{t}{\tau}\right)\exp\left(-\frac{l^2}{4\lambda^2 t/\tau}\right),$$

Where $\lambda$ is the length constant (characteristic length) and $\tau$ is the time constant of the axon.

Consider a Node of Ranvier (NoR) at $(x_0, y_0, 0)$ surrounded by myelinated axon on either side. At the node, the transmembrane current is $I_{AP}(t)$; based on this, we have

$$I(l,t) = -V_0/R_m + \delta(l)I_{AP}(t),$$

where $\delta$ is the delta distribution, $V_r$ is the resting membrane potential, and $V_r/R_m$ represents the leak current. By assuming that NoRs are maximally spatially separated [**Assumption 2.1a**], an AP will attenuate significantly before reaching the next node. From this, we make the critical approximation that $V_0(l) = V_r$; that is, prior to the activation at each node, the axonal membrane potential is at rest [**Assumption 2.1b**]. Given these assumptions, we have the equation

$$V(l,t) = V_{\rm r} e^{-t/\tau} + \frac{R_m}{\tau}\int_0^t \int_{-\infty}^{\infty} \big(-V_r/R_m + I_{AP}(t-s)\delta(l-x)\big) G(x,s)\,dx\,ds$$
$$= V_r + \frac{R_m}{\tau}\int_0^t \frac{I_{AP}(t-s)}{\sqrt{4\pi\lambda^2 s/\tau}} e^{-\frac{s}{\tau}}\, e^{-\frac{l^2}{4\lambda^2 s/\tau}} ds\ ,$$

and accordingly,

$$I_a(l,t) = -\frac{\pi d^2}{4R_a}\frac{\partial V}{\partial l} = \frac{\sqrt{\pi} d^2}{2R_a C_m}\int_0^t \frac{I_{AP}(t-s) l}{(4\lambda^2 s/\tau)^{3/2}} e^{-\frac{s}{\tau}} e^{-\frac{l^2}{4\lambda^2 s/\tau}} ds.$$

We can then compute the electrical field generated by an AP at a given NoR as

$$\phi_{AP}(t; x_0, y_0, \varphi, L)$$
$$= \frac{d^2}{8\sigma R_a C_m \pi^{3/2}}\int_0^t A(s)\Bigg[\int_{-\infty}^{L} \frac{(r_0-l) l e^{-\frac{l^2}{4\lambda^2 s/\tau}}}{(x_0^2+y_0^2+(r_0-l)^2)^{3/2}}\ dl$$
$$+ \int_L^{\infty} \frac{I_a(l,t)(x_0\cos\varphi + y_0\sin\varphi + l - L)}{(x_0^2+y_0^2+2x_0(l-L)\cos\varphi + 2y_0(l-L)\sin\varphi + (l-L)^2 + (r_0-L)^2)^{3/2}} l e^{-\frac{l^2}{4\lambda^2 s/\tau}}\ dl\Bigg]\, ds,$$

where $A(s) = I_{AP}(t-s)e^{-\frac{s}{\tau}}(4\lambda^2 s/\tau)^{-3/2}$. Our goal is then to compute the inner integrals.

Note that when $l^2 \gtrsim 36\lambda^2 s/\tau$, we have $e^{-\frac{l^2}{4\lambda^2 s/\tau}} \ll 1$ and $36\lambda^2 s/\tau \ll r_0$. Motivated by this observation, we let $\varepsilon > 1/r_0$ and define $N_\epsilon \ll r_0$ such that, for all $|l| < N_\varepsilon$, we have $e^{-\frac{l^2}{4\lambda^2 s/\tau}} < \varepsilon$. Note also that for all $|l| < N_\varepsilon$, we have $(r_0 - l) \approx r_0$. The following results thus hold for $r_0$ large; this is possible because the length constant along an axon is much less than the distance of the axon to the EEG electrode.

By *Assumption 2.1*, there is only one NoR within $N_\varepsilon$ length units of the axon bend ($L$). Thus, the contribution of an AP propagation along the entire axon can be described by the actions of this single NoR. For simplicity, we assume in what follows that this node is exactly on the bend, i.e. $L = 0$. However, the results can be generalized to the case where $L$ is arbitrary.

***First integral.***

$$\int_{-\infty}^{0} \frac{(r_0-l) l e^{-\frac{l^2}{4\lambda^2 s/\tau}}}{(x_0^2+y_0^2+(r_0-l)^2)^{3/2}}\ dl \approx \int_{-N_\epsilon}^{0} \frac{(r_0-l) l e^{-\frac{l^2}{4\lambda^2 s/\tau}}}{(x_0^2+y_0^2+(r_0-l)^2)^{3/2}}\ dl.$$

Since for $-N_\varepsilon < l \le 0$, we have $r_0 - l \approx l$, this integral can be solved as follows

$$\int_{-N_\epsilon}^{0} \frac{(r_0-l) l e^{-\frac{l^2}{4\lambda^2 s/\tau}}}{(x_0^2+y_0^2+(r_0-l)^2)^{3/2}}\ dl \approx \frac{r_0}{(x_0^2+y_0^2+r_0^2)^{3/2}}\int_{-\infty}^{0} l e^{-\frac{l^2}{4\lambda^2 s/\tau}}\ dl \approx \frac{2r_0\lambda^2 s/\tau}{(x_0^2+y_0^2+r_0^2)^{3/2}}$$

***Second integral.*** As above,

$$\int_0^\infty \frac{I_a(l,t)(x_0\cos\varphi + y_0\sin\varphi + l - L)}{(x_0^2+y_0^2+2x_0(l-L)\cos\varphi + 2y_0(l-L)\sin\varphi + (l-L)^2 + (r_0-L)^2)^{3/2}} l e^{-\frac{l^2}{4\lambda^2 s/\tau}}\ dl$$
$$\approx \int_0^{\mathrm{N}_\varepsilon} \frac{I_a(l,t)(x_0\cos\varphi + y_0\sin\varphi + l - L)}{(x_0^2+y_0^2+2x_0(l-L)\cos\varphi + 2y_0(l-L)\sin\varphi + (l-L)^2 + (r_0-L)^2)^{3/2}} l e^{-\frac{l^2}{4\lambda^2 s/\tau}}\ dl.$$

For $0 \le l < N_\varepsilon$ we have the following approximation

$$\frac{(x_0\cos\varphi + y_0\sin\varphi + l - L)}{(x_0^2+y_0^2+2x_0(l-L)\cos\varphi + 2y_0(l-L)\sin\varphi + l^2 + r_0^2)^{3/2}} \approx \frac{(x_0\cos\varphi + y_0\sin\varphi)}{(x_0^2+y_0^2+r_0^2)^{3/2}}.$$

That is, because $r_0 \gg N_\varepsilon$, the angle of the axon relative to the electrode is approximately constant over the distance that the AP decays. This gives us the solution

$$\int_0^{N_\epsilon} \frac{(x_0\cos\varphi + y_0\sin\varphi)}{(x_0^2+y_0^2+r_0^2)^{3/2}} l e^{-\frac{l^2}{4\lambda^2 s/\tau}}\, dl \approx \frac{2(x_0\cos\varphi + y_0\sin\varphi)\lambda^2 s/\tau}{(x_0^2+y_0^2+r_0^2)^{3/2}}$$

Combining these two results, we arrive at the solution

$$\phi_{AP}(t;x_0,y_0,\varphi,L) = \frac{d^2(x_0\cos\varphi + y_0\sin\varphi + r_0)}{32\lambda\sigma R_a C_m (x_0^2+y_0^2+r_0^2)^{3/2}\pi^{3/2}} \int_0^t \frac{I_{AP}(t-s)}{\sqrt{s/\tau}} e^{-\frac{s}{\tau}} ds$$

Again, this equation gives us an approximate response of the extracellular electric field to an AP propagating along an axon.

Typically, AP currents are modelled using a Hodgkin-Huxley type formalism with voltage-gated potassium and sodium channels. To allow for an analytical solution, what we do instead here is explicitly define the current as follows

$$I_{AP}(t) = \frac{\gamma_{AP}(Be^{-t/\tau_1} - e^{-t/\tau_2})}{(1-\tau_1/\tau_2)(B\tau_1/\tau_2)^{-\frac{\tau_1}{\tau_1-\tau_2}}}$$

which provides a physiologically reasonable approximation for the AP waveform (Figure S3A). By using this current and letting

$$\tilde{A}(x_0,y_0,\varphi) := \frac{d^2(x_0\cos\varphi + y_0\sin\varphi + r_0)}{32\lambda\sigma R_a C_m (x_0^2+y_0^2+r_0^2)^{3/2}\pi^{3/2}} \qquad (L=0),$$

we obtain the following solution

$$\begin{aligned}
\phi_{AP}(t;x_0,y_0,\varphi,0) &= \tilde{A}(x_0,y_0,\varphi) \int_0^t \frac{I_{AP}(t-s)}{\sqrt{s/\tau}} e^{-\frac{s}{\tau}} ds \\
&= \tilde{A}(x_0,y_0,\varphi)\gamma_{AP}\sqrt{\tau}\left[\int_0^t \frac{Be^{-(t-s)/\tau_1}e^{-s/\tau}}{\sqrt{s}} ds - \int_0^t \frac{e^{-(t-s)/\tau_2}e^{-s/\tau}}{\sqrt{s}} ds\right] \\
&= \tilde{A}(x_0,y_0,\varphi)\sqrt{\pi\tau}\gamma_{AP}\left[\frac{Be^{-t/\tau_1}}{\sqrt{1/\tau_1 - 1/\tau}} \operatorname{erf}\left(\sqrt{t\left(\frac{1}{\tau_1} - \frac{1}{\tau}\right)}\right)\right. \\
&\qquad \left. - \frac{e^{-t/\tau_2}}{\sqrt{1/\tau_2 - 1/\tau}} \operatorname{erf}\left(\sqrt{t\left(\frac{1}{\tau_2} - \frac{1}{\tau}\right)}\right)\right].
\end{aligned}$$

Accordingly, we define

$$\alpha_{AP}(t)$$

$$:= \frac{\gamma_{AP}}{\lambda \mathrm{C_m}} \begin{cases} \frac{B\sqrt{\tau}e^{-t/\tau_1}}{\sqrt{1/\tau_1 - 1/\tau}} \operatorname{erf}\left(\sqrt{t\left(\frac{1}{\tau_1} - \frac{1}{\tau}\right)}\right) - \frac{\sqrt{\tau}e^{-t/\tau_2}}{\sqrt{1/\tau_2 - 1/\tau}} \operatorname{erf}\left(\sqrt{t\left(\frac{1}{\tau_2} - \frac{1}{\tau}\right)}\right), & t \geq 0 \\ 0, & t < 0 \end{cases}$$

whose graph is displayed in Figure S3C. As in the case for EPSPs and IPSPs, we treat $\alpha_{AP}$ as the impulse response of the electric field to an AP and write the electric field generated by a single axon as

$$\phi^i_{AP}(t) = A(x_i, y_i, \varphi_i)(\alpha_{AP} \star \tilde{w}_i)(t),$$

where $\tilde{w}_i(t)$ is a Poisson point processes with a mean rate $\lambda_{AP}$ (representing the occurrence of APs) and $A = \sqrt{\pi}\tilde{A}$. The total contribution from APs is then computed by summing over all such axons

$$\phi_{AP}(t) = \sum_i \phi^i_{AP}(t) = \int_0^\infty d\tau \ \alpha_{AP}(t-\tau) \sum_i A(x_i, y_i, \varphi_i)\tilde{w}_i(\tau).$$

As done previously, we assume that the cortex is a continuum of axons with a uniform density, $\rho_{AP}$. Furthermore, we assume that $\varphi_i \sim \mathcal{U}_{[0,2\pi]}$ is uniformly distributed. This gives us

$$\phi_{AP}(t) = \int_0^\infty d\tau \ \alpha_{AP}(t-\tau) \int_{-\infty}^\infty dx \int_{-\infty}^\infty dy \int_0^{2\pi} d\varphi A(x, y, \varphi) w_i(\tau)$$

where $w_i(\tau)$ is a Poisson process with a mean rate $\lambda_{AP}\rho_{AP}/2\pi$ . As in the case of synaptic activity, we rewrite this as

$$\phi_{AP}(t) = (\alpha_{AP} \star W_{AP})(t),$$

where

$$\sigma^2_{AP} = \int_0^\infty dr \int_0^{2\pi} d\theta \int_0^{2\pi} d\varphi \, A(r,\theta,\varphi)^2 \, \frac{r\rho_{AP}\lambda_{AP}}{2\pi}$$
$$= \frac{d^4\lambda_{AP}\rho_{AP}}{2048\sigma^2 R_a^2 C_m^2 \lambda^2 \pi^3} \int_0^\infty \int_0^{2\pi} \int_0^{2\pi} drd\theta d\varphi \frac{(r \, cos\, \theta \, cos\, \varphi + r \, sin\, \theta \, sin\, \varphi + r_0)^2 r}{(r^2 + r_0^2)^3}$$
$$= \frac{d^4\lambda_{AP}\rho_{AP}}{768\sigma^2 R_a^2 r_0^2 \pi}.$$

**Note S3. Consequences of synchronous neural activity on EEG power spectrum.**

In order to include oscillatory network dynamics, we model the trains of EPSPs ($w_E$), IPSPs ($w_I$), and APs ($w_{AP}$) as cyclic Poisson processes with rate functions of the form $\Lambda(t;\lambda_x) = N_x\lambda_x(1 + c\,sin(2\pi k_t t))$, for $x = E, I$, or $AP$, where $k(t)$ is the frequency of oscillation that can vary with time, and $c < 1$ combines the fraction of synapses whose inputs are synchronized and the degree to which this synchronization modulates the base firing rate.

Due to Assumption 1.1 (radial symmetry), if synaptic inputs are synchronized, then their electric fields should cancel out, not just on average, but at every moment of time, leading to no net contribution to the EEG. We therefore introduce an asymmetry coefficient, $a$. Physiologically, this coefficient captures the asymmetric propagation of synaptic currents in pyramidal cells, whereby, due to the apical tufts, there is thought to be a net dipole orientation along the apical-basal axis[7]. The implication of this asymmetry is precisely that, given synchronous activity, only the dipoles from $(1-a)cN_x$ synapses will cancel out, leaving us with the following equation for the rate of EEG-contributing dipole moments

$$\Lambda(t;\lambda_x, b, a) = N_x\lambda_x[1 - c + a \cdot c + a \cdot c\sin(2\pi k_t t)]$$

If $k(t) = k_0$ is constant and $a \cdot c \ll 1$, then

$$\mathbb{E}|\hat{w}_x(f)|^2 \approx N_x\lambda_x(1 - \mathrm{c} + \mathrm{ac}) + (acN_x\lambda_x)^2\delta(f - k_0).$$

If, on the other hand, $k(t)$ fluctuates slowly, we can approximate $\hat{w}_x$ as

$$\mathbb{E}|\hat{w}_x(f)|^2 \approx N_x\lambda_x(1 - \mathrm{c} + \mathrm{ac}) + (acN_x\lambda_x)^2 g_k(f),$$

where $g_k(f)$ is a function representing the distribution of frequencies that the rhythm adopts over a given window of time, e.g. Figure 2G.

We note that simulating a power spectrum containing peaks with realistic amplitudes is achievable with $a \approx 0.01$ and $c \approx 0.01$ (see figure below), suggesting that our assumption $a \cdot c \ll 1$ is reasonable.

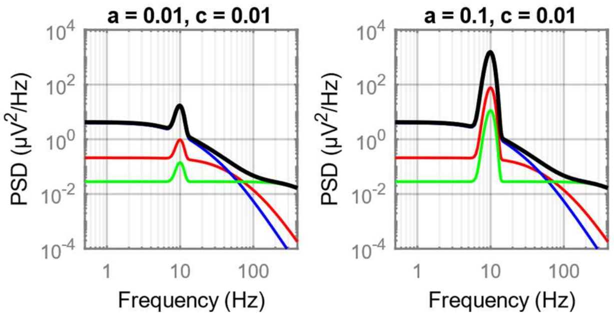


This observation also validates Assumption 1.1, i.e. that during asynchronous activity, any net asymmetry in dipole orientation is negligible and may be ignored. Finally, since we expect $a$ and $c$ to be in general small, we let $N_x\lambda_x(1 - c + ac) \approx N_x\lambda_x$ and $b \coloneqq N_x\lambda_x ac$, leading to the following equation for the expected power spectrum of the EEG

$$\mathbb{E}|\hat{\phi}_{EEG}(f)|^2 = \Lambda_{\mathrm{E}}(1+bg_k(f))|\hat{\alpha}_E|^2 + \Lambda_{\mathrm{I}}(1+bg_k(f))|\hat{\alpha}_I|^2 + \Lambda_{AP}(1+bg_k(f))|\hat{\alpha}_{AP}|^2$$

where, for example, $g_k(f)$ would be centred around 10 Hz for alpha rhythms. This equation extends linearly for multiple modulatory rhythms.
