## Supplemental Figures for "Aperiodic EEG activity masks the dynamics of neural oscillations during loss of consciousness from propofol"

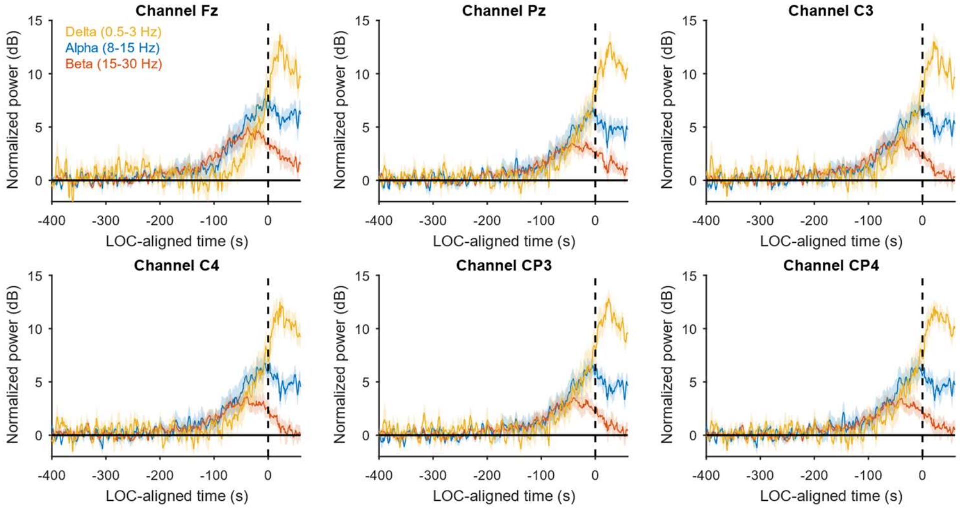


**Figure S1. Band power changes at different EEG recording locations. Related to Figure 1**.

Same as Figure 1C, but for other electrode locations, i.e. Fz, Pz, C3, C4, CP3, CP4.

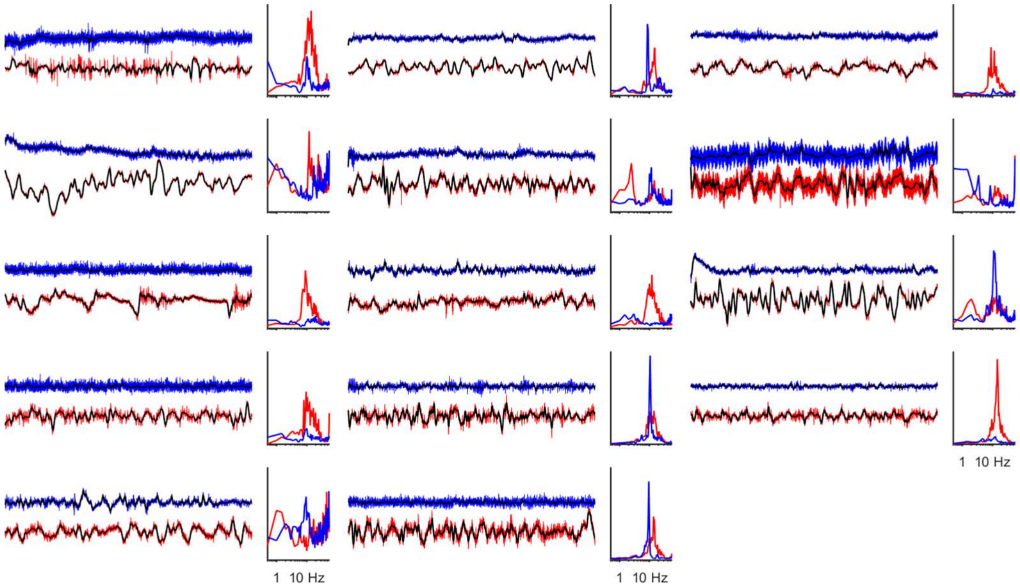


**Figure S2. Emergence of slow rhythms following LOC. Related to Figure 2.** All time series shown are 25 s long. Blue traces are examples of baseline EEG signals for each of the 14 patients, and in red are example traces post-LOC. Superimposed black lines indicate results of low pass filter of the time series, as in Figure 2. To the right of each pair of time series is the 1/f detrended power spectrum at baseline (blue) and post-LOC (red). The appearance of slow oscillations in many of the time series is not apparent in the power spectra after 1/f detrending, indicating that the 1/f fitting is removing low frequency rhythmic power.

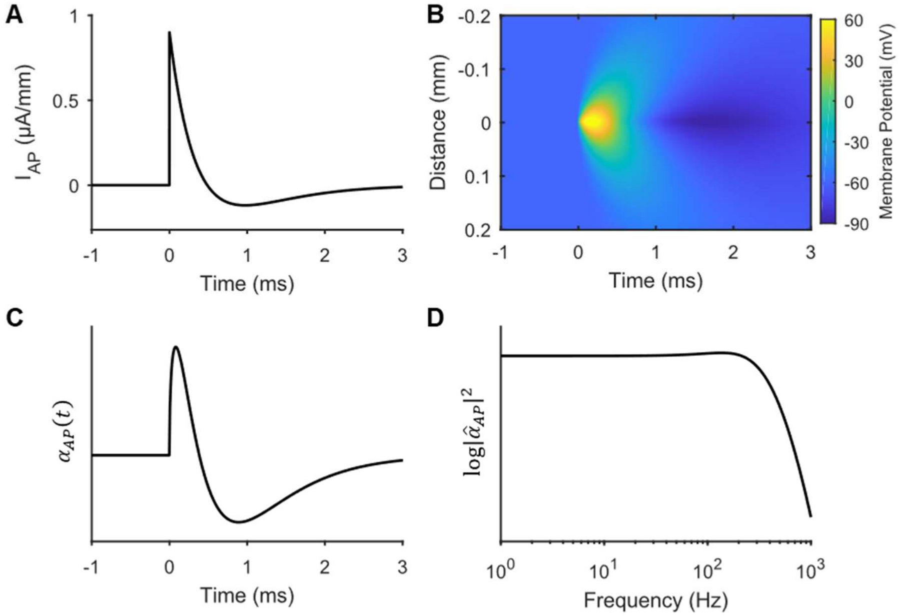


**Figure S3**. **Contributions of APs to extracellular field. Related to Figure 3.**

**(A)** Time course of the modelled current at a Node of Ranvier (NoR), $I_{AP}(t) = \gamma_{AP}(0.67e^{-t/0.6} - e^{-t/0.4})$ for $t \geq 0$.

**(B)** Simulation of the membrane potential along an axon given the following input current $I(t, x) = I_{AP}(t)\delta(x)$, with all other parameters from Table 1.

**(C)** Plot of the function $\alpha_{AP}(t)$.

**(D)** Numerically computed power spectrum of $\alpha_{AP}(t)$.

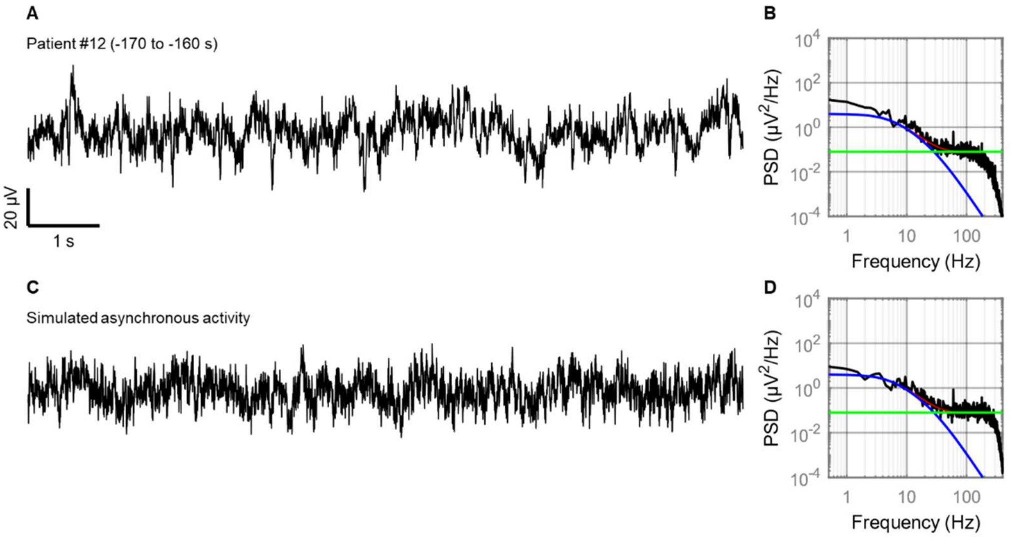


**Figure S4**. **Example of EEG with little sustained rhythmic activity. Related to Figure 3.**

**(A)** 10 s sample of EEG from patient #12, 170 s before LOC (45 s before propofol infusion).

**(B)** Same as in Figure 3C.

**(C)** Stochastic simulation of EEG model using scaling factors and time constants of IPSPs and APs estimated from panel B, bandpass filtered between 0.1-300 Hz.

**(D)** Power spectrum of the model simulation in panel C, superimposed with the analytical power spectrum of the underlying model.

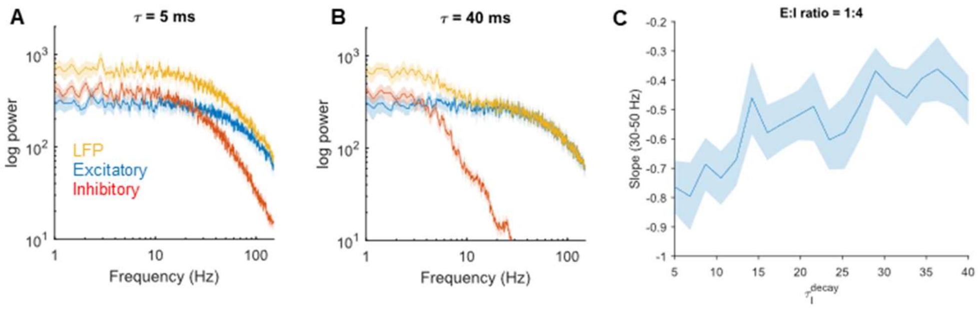


**Figure S5. Effect of inhibitory decay time on spectral slope. Related to Figure 4.**

**(A)** The computational model from Gao et al. (2017) was simulated with an excitatory/inhibitory (E:I) ratio of 1:4; coloured lines and shading represent mean ± S.E across 20 simulations. All parameters values were taken from Table 1 of Gao et al. (2017), except for the decay time of $GABA_A$ which was set here to 5 ms.

**(B)** Same as in panel A, but the decay time of $GABA_A$ is set to 40 ms. The frequency of inhibitory events was reduced to maintain an E:I ratio of 1:4.

**(C)** As the decay time of inhibition is increased, while maintaining a constant E:I ratio, the slope of the power spectrum between 30-50 Hz increases.
